# A theory of multi-task computation and task selection

**DOI:** 10.64898/2025.12.12.693832

**Authors:** Owen Marschall, David G. Clark, Ashok Litwin-Kumar

## Abstract

Neural activity during the performance of a stereotyped behavioral task is often described as low-dimensional, occupying only a limited region in the space of all firing-rate patterns. This region has been referred to as the “neural manifold” associated with a task. More recently, recordings of neural activity in animals challenged to perform multiple tasks have suggested that each task is associated with a different low-dimensional manifold. What connectivity structures underlie this flexibility in neural dynamics, and how is interference between the dynamics associated with different tasks avoided? We develop a theoretical model for multi-task computation in nonlinear recurrent neural networks whose connectivity is constructed as a weighted sum of many low-rank components, each encoding the dynamics associated with a different task. The model demonstrates that interference between different tasks’ dynamics limits flexible multi-tasking and can lead to chaotic fluctuations. However, small modulations of a network’s effective connectivity overcome this interference. We derive the conditions that enable such task selection and characterize both single-neuron and population statistics in task-selected and unselected states. The model reveals the requirements for a single network to produce distinct dynamics confined to distinct neural manifolds and suggests circuit mechanisms that support this capability. Using the model, we propose different hypotheses for explaining the origin of high-dimensional neural activity in large-scale recordings.

## Introduction

Adaptive behavior requires organisms to perform different neural computations at different times or in different contexts. For example, both spontaneous and skilled, practiced movements involve sequencing discrete motor computations [1, 2]. Similarly, cognitive tasks require operations on task-relevant variables that may differ across task, condition, or epoch [3–7]. Numerous studies support the proposal that individual neural computations are associated with coordinated dynamics of interconnected populations of neurons [8], implying that the effective dynamical system governing these neurons’ activity must rapidly change when a new computation is to be performed. Little is known about the circuit mechanisms that underlie these dynamical switches.

During execution of a single task, population activity is typically confined to a low-dimensional neural manifold [9, 10]. Mixed selectivity—the observation that individual neurons often participate in multiple distinct tasks [3, 5, 11, 12], although not always [13]—implies that these manifolds are distributed across many neurons rather than aligned to a small subset. This distributed structure is reproduced in recurrent neural networks trained to perform multiple tasks [4, 14]. Recent work shows that the location and orientation of the manifold can change depending on the task being performed [2, 15–18]. Thus, switching between manifolds, each distributed across the population, may underlie the rapid changes in neural dynamics and behavior required for switching tasks [2].

In another line of work, neural activity has been observed to be high-dimensional during periods of spontaneous, uninstructed behavior [19, 20], in contrast to observations of low-dimensional activity during the performance of stereotyped tasks [21, 22]. These observations may be compatible with the network switching among many distinct manifolds, each associated with a different self-initiated behavior [1, 23]. The union of many such individually low-dimensional manifolds can be high-dimensional if they are heterogeneously oriented in neural state space, as recent experimental work suggests [2]. Alternatively, such activity may reflect unstructured, high-dimensional fluctuations, as proposed by random-network models of the spontaneous state [24, 25].

Altogether, these observations raise several theoretical questions. What forms of recurrent connectivity support many distinct task-related manifolds, with mixed selectivity to different tasks at the single-neuron level? Under what conditions do the dynamics associated with each manifold interfere or not interfere? How does a circuit selectively engage the dynamics appropriate for one task while suppressing others? What conditions lead to high-dimensional neural activity?

Prior work has addressed multi-tasking by training recurrent neural networks on multiple tasks using machine learning methods and then analyzing the solutions [3, 4, 14, 26, 27]. Such studies have shown that artificial networks reuse dynamical motifs, such as fixed points and ring attractors, when the tasks have shared component subtasks. We take a complementary approach, using a solvable network model rather than trained networks. For simplicity, we assume each “task” is an irreducible computation that can be solved by a single low-dimensional dynamical system, without compositional overlap [6].

Specifically, we propose and analyze a mathematically tractable model of multi-tasking neural circuits, in which distinct task-related dynamics operate on distinct low-dimensional activity manifolds. Building on low-rank network models that explain how recurrent structure shapes low-dimensional dynamics [28, 29], we model networks whose weight matrix is constructed as a sum of many low-rank single-task matrices [30]. Because the network dynamics are nonlinear, this leads to complex interactions between the dynamics associated with different tasks. We develop a dynamical mean-field theory that characterizes how tasks interfere, how spontaneous fluctuations arise when a system has learned to perform many distinct tasks, and how these dynamics manifest at both the single-neuron and population levels. Our model makes predictions about single-neuron tuning and fluctuations, trial-to-trial variability in neural activity projected onto task-related dimensions, and the dimensionality of population activity that can be examined in experimental data.

A key feature of the model is that small modulations of connectivity can lead to the dynamic selection of a single task and the suppression of others. This provides a biologically plausible mechanism for task selection. In particular, our proposal is consistent with evidence that external inputs, for example thalamic inputs, gate and sustain motor cortical activity [31, 32]. More generally, task execution may result from coordination between recurrent dynamics and modulatory pathways. In total, we provide a theory relating connectivity, dynamics, and experimentally measurable population signatures of multi-tasking in recurrent neural circuits.

## Results

We study nonlinear recurrent neural networks of *N* continuous-time rate neurons with currents *x*_*i*_(*t*) and firing rates *ϕ*_*i*_(*t*) that evolve according to

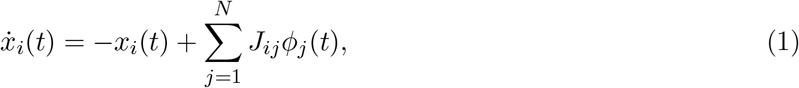

with a sigmoid-shaped firing-rate function (taken to be 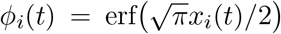 in simulations). The network dynamics are controlled by the *N* × *N* weight matrix **J** with elements *J*_*ij*_. We assume that *N* is large, and our theoretical results are exact in the *N* → ∞ limit. We now discuss the dynamics that occur when **J** is low-rank, before presenting our model in which many such low-rank matrices are superposed.

### Low-rank networks

Many previous works have considered the case where the recurrent connectivity **J** is low-rank [28, 29, 33, 34], given by

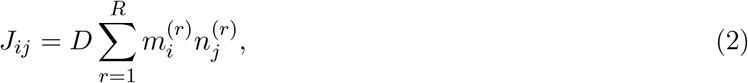

a sum over a small number *R* ≪ *N* of outer products of *N*-dimensional *loading vectors* **m**^(*r*)^, **n**^(*r*)^, scaled by a constant *D* ≥ 0. The right loading vectors **n**^(*r*)^ define the subspace of neural firing rates that determines the recurrent input to the network, and this input lies in a subspace spanned by the left loading vectors **m**^(*r*)^ (Fig. 1A, left). When transformed by the firing-rate nonlinearity, this *R*-dimensional input subspace produces neural activity that lies on an *R*-dimensional nonlinear manifold (Fig. 1A, right).

**Figure 1:**
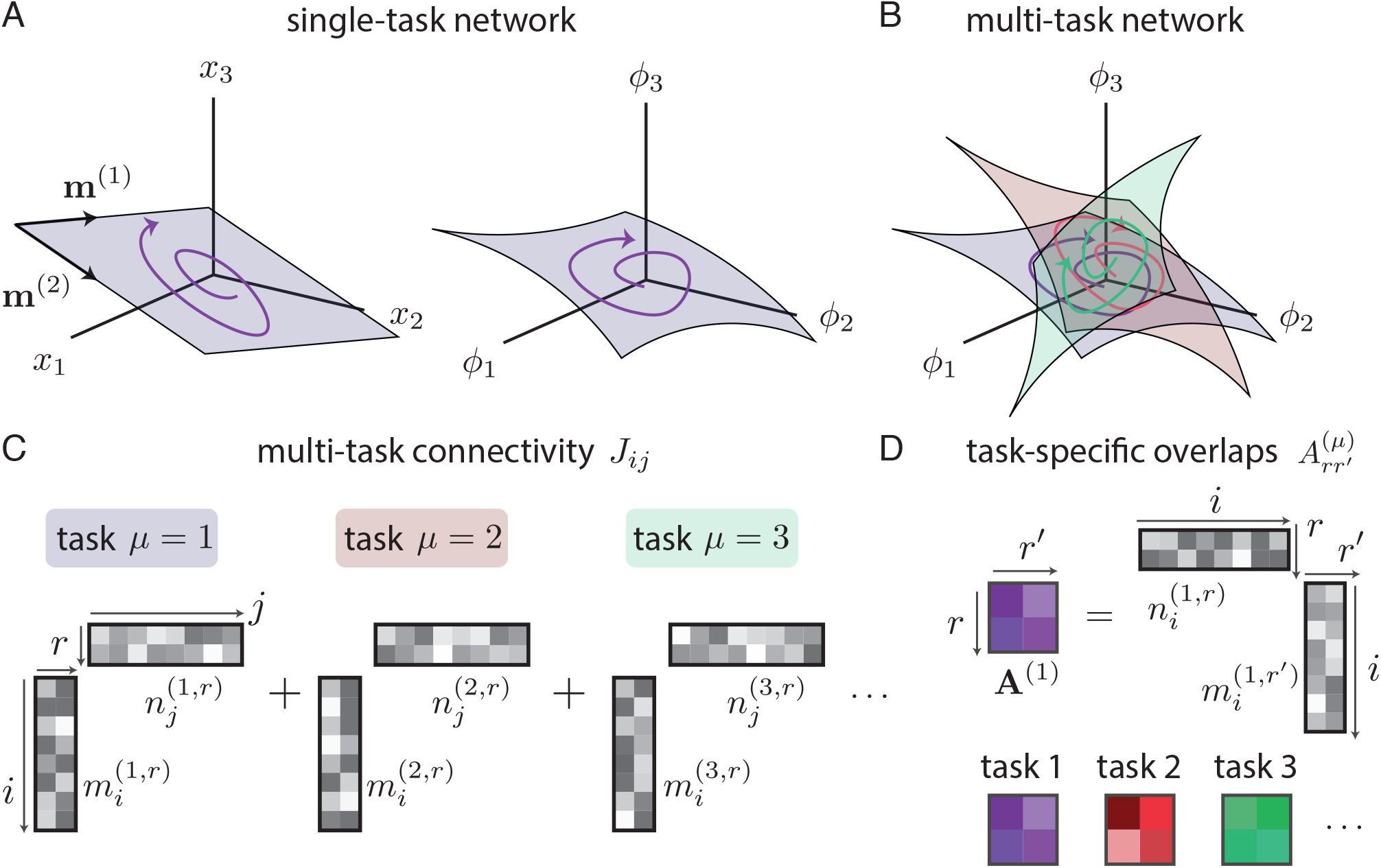
Single-task and multi-task network dynamics. **(A)** Left: Illustration of a state-space trajectory in the space of neuronal pre-activations (purple), confined to a task-relevant two-dimensional subspace. Such a geometry arises in rank-two networks with loading vectors **m**^(1)^, **m**^(2)^. Right: Illustration of the same trajectory in firing-rate space. Activity lies on a two-dimensional nonlinear manifold. **(B)** Schematic of a “multi-task” network that flexibly implements multiple low-dimensional dynamics. Each task is associated with a different nonlinear manifold, which collectively span more activity dimensions than any one manifold. **(C)** Schematic of multi-task network connectivity as a sum of low-rank matrices, each associated with a different task. **(D)** Each task is associated with an overlap matrix comprising pairwise inner products among the left and right loading vectors.

We consider the special case of Gaussian-distributed loading-vector components with zero mean and specified covariance structure. All components are generated i.i.d. across different neurons. For a given neuron *i*, the left- and right-loading-vector components are sampled with nonzero covariances 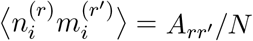 according to the entries of the *R* × *R overlap matrix* **A**. As such, this matrix encodes the loading vectors’ geometrical arrangement 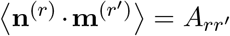 and plays a major role in determining the network’s dynamics.

Activity in a low-rank network can be described by a set of *latent variables*, defined by projections onto the right loading vectors, *z*_*r*_(*t*) = *D*(**n**^(*r*)^ *·* ***ϕ***(*t*)). In the *N* → ∞ limit, the overlap matrix determines the dynamics among the latent variables according to

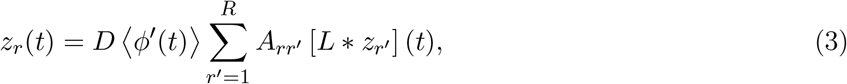

where *L*(*τ*) is a low-pass filter with the single-neuron time constant (see Methods). The gain factor ⟨*ϕ*′(*t*)⟩ is the population-averaged slope of the firing-rate nonlinearity. It depends on the norm of the low-pass-filtered latent variables, with larger norm reducing the gain due to the sigmoidal form of *ϕ*(·) (see Methods). Eq. (3) summarizes a low-rank network’s activity as an *R*-dimensional, nonlinear dynamical system [28, 29]. The gain factor ⟨*ϕ*′(*t*)⟩ is the only nonlinearity in this system and thus is necessary for nonlinear dynamical behaviors such as stable limit cycles or multistability.

### Multi-task low-rank networks

Neural recordings suggest that the activity of individual circuits can switch among distinct neural manifolds that collectively span a large number of dimensions [Fig. 1B; 2, 16]. When measured across many different contexts, such activity is high-dimensional, as are measurements of dimension during spontaneous behavior [19, 20]. Such phenomena are not well modeled by low-rank networks that produce activity residing on a single low-dimensional manifold. We therefore developed a model of multi-task networks in which distinct dynamics occur on distinct neural manifolds.

In the context of our model, “task” refers to the low-dimensional dynamics associated with a particular manifold in firing-rate space, which we interpret as implementing a specific computation. Indeed, prior work in the single-task setting has shown that low-rank networks can implement experimentally inspired input–output mappings through their low-dimensional dynamics [28, 34]. Although real tasks involve both sensory inputs and behavioral outputs, we focus only on the recurrent dynamics, noting that our model could be easily extended to include task-specific inputs as in prior single-task works (see Discussion).

The network performs *P* different tasks, indexed by *µ* = 1, …, *P*. Each task is associated with a rank-*R* matrix of the form of Eq. (2), and the multi-task weight matrix is constructed as a weighted sum of these matrices:

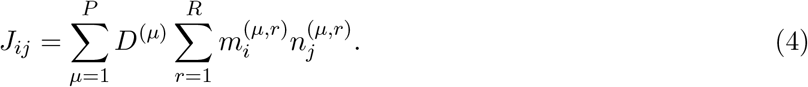

Each task *µ* is therefore parameterized by *R* left and right loading vectors **m**^(*µ,r*)^ and **n**^(*µ,r*)^, and a weight *D*^(*µ*)^ ≥ 0 controlling the strength of the connectivity component associated with that task. The loading vectors are Gaussian and i.i.d. across neurons, as in the single-task setting. They are furthermore independent across tasks, i.e. there is zero expected overlap between loading vectors of distinct tasks. The covariance for task *µ* given by

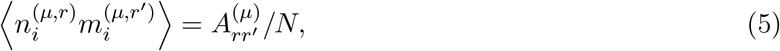

with the quantities 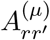 comprising the task-specific *R*×*R* overlap matrix **A**^(*µ*)^ (Fig. 1D). We refer to the subspace spanned by **m**^(*µ,r*)^ for *r* = 1, …, *R* as *task subspace µ*, the *R*-dimensional space that contains the input to the network associated with task *µ*.

As described previously for single-task networks, the structure of overlaps between left and right loading vectors plays a major role in shaping network dynamics. The connectivity of Eq. (4) can be interpreted as that of a rank-*P R* network with a particular structure of overlaps among left and right loading vectors. Specifically, the overlaps between **m**^(*µ,r*)^ and **n**^(*ν,r′*)^ are constrained to be, on average, block-diagonal, with the block entries given by 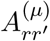 (Fig. S1). Each of these blocks corresponds to *R*-dimensional dynamics associated with a single task subspace. Our model therefore describes synaptic weight matrices with strong, structured interactions among the components associated with a single task, specified by these blocks, and small, random overlap between components associated with different tasks. In terms of activity, the subspaces corresponding to different tasks are nearly, although not exactly, orthogonal when *N* is large, with finite-size fluctuations in cross-task overlaps that are reduced as *N* grows. This blockwise constraint reduces the number of parameters needed to specify the model from *P*^2^*R*^2^ to *PR*^2^ overlap-matrix elements—a dramatic reduction when *P* is large—while allowing the possibility of distinct tasks occupying different manifolds defined by the connectivity.

### Multi-task networks exhibit winner-take-all competition across tasks

We begin by comparing single-task networks (*P* = 1) to a minimal multi-task network (*P* = 2). Network weights are set according to Eq. (4), fixing the within-task dimension *R* = 2. The *P* = 1 case reduces to the low-rank networks described by Eqs. (2) and (3). Such low-rank networks can produce a variety of different dynamics. For example, two different choices of a 2 × 2 overlap matrix **A** produce networks that exhibit either a stable limit cycle (Fig. 2A) or bistability (Fig. 2B). We next consider a multi-task network with *P* = 2 using a weight matrix that superposes the two matrices used above that, on their own, produce limit-cycle and fixed-point dynamics. The dynamics can be analyzed via two sets (one for each task) of latent variables 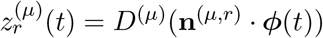 where *µ* = 1, 2 indexes tasks and *r* = 1, 2 indexes within-task dimensions. Both sets of latent variables initially follow trajectories similar to those of the corresponding single-task networks (Fig. 2C). However, closer examination reveals that the dynamics are coupled across tasks and exhibit different asymptotic behaviors. While the dynamics in the task-1 (limit cycle) subspace are barely changed (Fig. 2A vs. Fig. 2C, left), the dynamics in the task-2 (bistable attractor) subspace lose their bistability. Trajectories are initially attracted towards one of the two previously stable fixed points but subsequently decay to the origin, once task-1 activity has grown sufficiently large in magnitude (Fig. 2B vs. Fig. 2C, right). The network therefore exhibits winner-take-all behavior, where one task dominates and suppresses others. In this example, task 1 dominates because *D*^(1)^ > *D*^(2)^. Reversing this inequality causes task 2 to dominate (Fig. S2).

**Figure 2:**
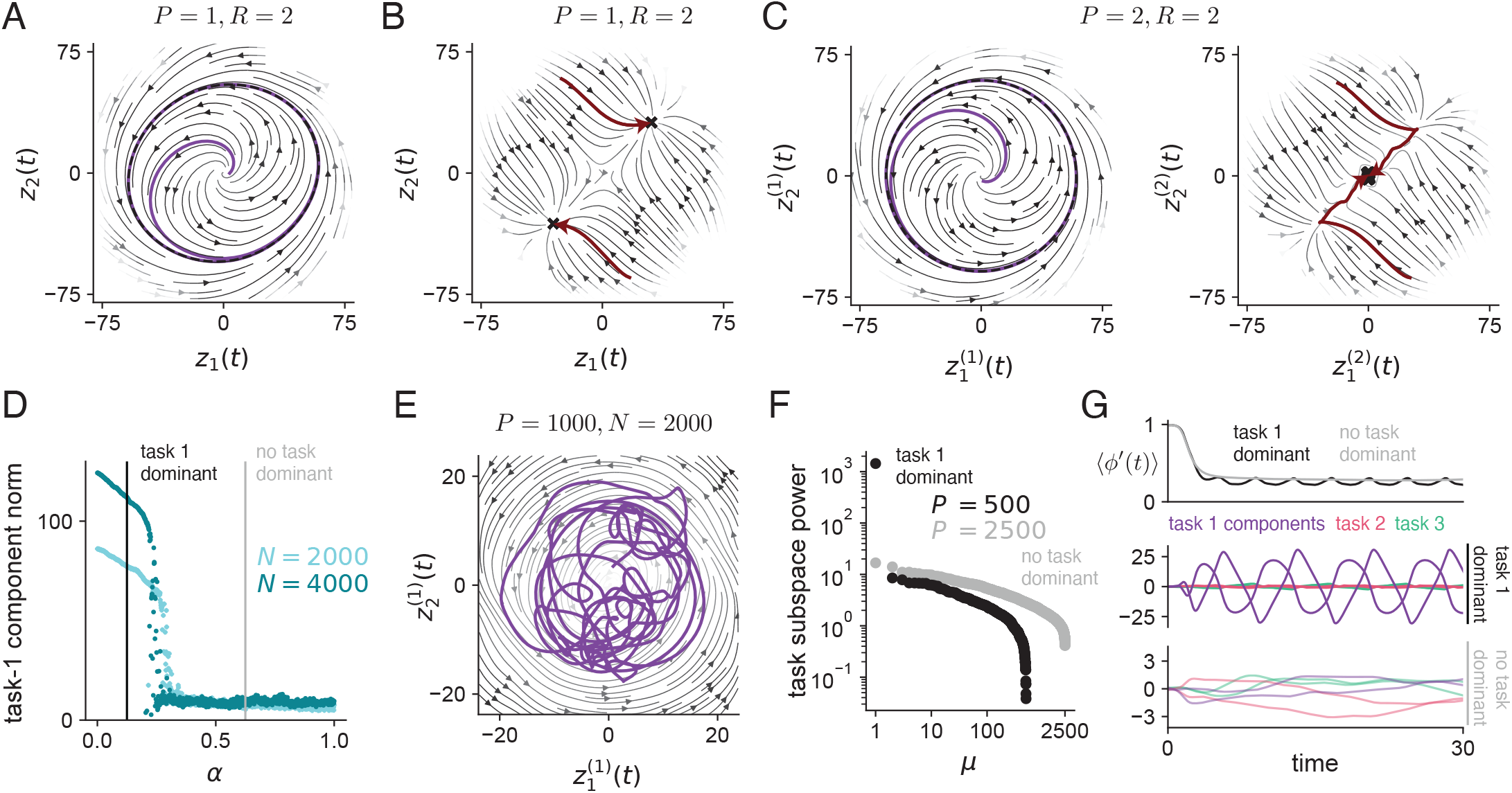
Dynamics of networks with varying *P*. (See Methods for parameter values.) **(A)** Latent-variable flow fields (black) and a latent-variable trajectory (purple) in a single-task network with **A** chosen to produce limit-cycle dynamics. Dashed line: limit cycle. **(B)** Latent-variable flow fields (black) and latent variable trajectories for two separate runs with different initial conditions (red) in a single-task network with **A** chosen to produce two stable fixed points. Black X’s: final time points for all trajectories used to generate flow fields. **(C)** A two-task network with *D*^(1)^ > *D*^(2)^. Left: flow fields (black) of projections 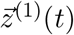 onto task-1 subspace and a latent-variable trajectory (purple). Right: flow fields (black) of projections 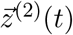 onto task-2 subspace simulated over many initial conditions, as well as two example trajectories (red). Black X’s: final time points for all trajectories. **(D)** Time-averaged latent variable norm 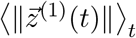 for task *µ* = 1 as a function of the number of total tasks *α* = *P*/*N* normalized by *N*. Same overlap parameters as (C) for *µ* = 1, 2, and beyond *µ* > 2 we randomly sampled 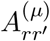. **(E)** Same plot of task-1 dynamics as in left panel of (C) but for a network with *P* = 1000 and smaller axis scales. **(F)** Time-averaged power 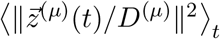 of the projections onto all task subspaces, normalized by *D*^(*µ*)^ and sorted in descending order, for a network with *P* = 500 tasks (black) and a network with *P* = 2500 tasks (gray) from panel (D). **(G)** Top: average gain as a function of time for two different networks, one with no task dominant (gray) and one with task 1 dominating (black). Middle: normalized latent-variable trajectories 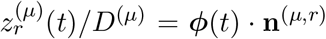 for each task in the task-1-dominant network. Bottom: same as middle for the network with no task dominant; note the difference in y-axis scale.

In a linear network, activity in distinct task subspaces evolves independently when the loading vectors for different tasks are orthogonal. Interference between tasks, defined as the influence of one task’s latent variables on another’s, occurs only when this orthogonality is broken. In multi-task networks with a fixed number of tasks, such interference becomes negligible as *N* → ∞, because randomly oriented activity patterns in a high-dimensional activity space are nearly orthogonal with high probability. What, then, is the source of the winner-take-all interference observed in Fig. 2C? In nonlinear networks with finite *P*, Eq. (3) still describes each task’s dynamics, but the gain factor now depends on the dynamics associated with all tasks [29]. Increased activity in any task subspace reduces the gain factor and suppresses the dynamics in other subspaces (see Methods). Consequently, at the steady state, the strongest task’s latent variables dominate and evolve as they would in a single-task network, with all others suppressed. This form of interference persists even in the limit *N* → ∞ and is unique to nonlinear networks that exhibit mixed selectivity and thus share neurons across tasks. In the example of Fig. 2C, the growth of task 1’s oscillatory latent variables suppresses dynamics in the task-2 subspace and causes trajectories to approach the origin, eliminating bistability.

These properties are not specific to networks with our particular choice of nonlinearity, as we observe similar winner-take-all behavior for other saturating nonlinearities, as well as in networks with non-saturating nonlinearities that are stabilized through recurrent inhibition [35] (Fig. S3). However, the fact that at most one task can dominate, rather than multiple tasks dominating simultaneously, is a consequence of Gaussian-distributed loadings, a simplification we use for mathematical tractability (see Discussion).

### Chaotic fluctuations in networks with large *P*

Higher cortical areas participate in many distinct tasks, so our primary focus is on networks with large *P*. We therefore simulate networks with increasing *P*. We configure the overlap matrix of task 1 to produce a stable limit cycle as before, while the overlap matrices of other tasks are randomly generated. We also set *D*^(1)^ to be large enough that task 1 would dominate any other task in isolation.

As *P* increases, the average magnitude of the projection of neural activity onto the task-1 subspace first decreases gradually, then drops sharply (Fig. 2D). This drop corresponds to a transition from a state in which task 1 is dominant and exhibits a stable limit cycle to a state in which no task is dominant. The point at which this transition occurs corresponds to a fixed ratio of number of tasks to number of neurons, denoted by *α* = *P*/*N* (Fig. 2D, light vs. dark cyan).

In an example network with large *P* and no task dominant, the task-1 latent variables fluctuate near the origin and are smaller in magnitude than those of networks with smaller *P* (Fig. 2E, purple trajectory). However, signatures of the task’s oscillatory structure remain visible in an average flow field computed over many samples of network activity (Fig. 2E, black). As we will show later, the irregular trajectories of latent variables in networks with large *P* are driven by chaotic fluctuations that arise from many small overlaps among distinct tasks’ subspaces, an effect that is vanishing for small *P* but significant for large *P*. In this state, activity is distributed across task subspaces (Fig. 2F, gray), rather than residing primarily in a dominant-task subspace (Fig. 2F, black). Additionally, the gain factor equilibrates to a time-independent value (Fig. 2G top, gray), rather than being dynamically entrained to any one task’s latent variables (Fig. 2G top, black).

The simulations shown in Fig. 2 demonstrate that nonlinear multi-task networks exhibit interference. When *P* is small, interference leads to winner-take-all dynamics in which one fixed task always dominates, regardless of the initial condition. As *P* is increased, interference additionally leads to chaotic fluctuations that eventually suppress task-related dynamics. Unlike previous models in which chaos arises due to unstructured, random connectivity [36], or weight matrices with both unstructured and low-rank components [28], in our model all components of the synaptic weight matrix are task-related (Eq. 4). This suggests that chaotic fluctuations, which are often proposed as an explanation for irregular neural activity recorded *in vivo*, can arise solely as a consequence of learning many tasks (see Discussion).

### Modulation of effective connectivity enables task selection

The results so far show that multi-task networks with fixed weights of the form of Eq. 4 cannot flexibly transition between distinct task-related activity patterns. Instead, at most one task dominates the winner-take-all dynamics (Fig. 2). We investigated mechanisms to circumvent this limitation. One possibility is to provide an input aligned with the task subspace of the desired task. Surprisingly, for networks with Gaussian loading vectors, this strategy does not enable task selection (Fig. S4) because such inputs modulate the gain factor by an equal amount across all tasks [29].

We therefore hypothesize that a mechanism that modulates *D*^(*µ*)^, the strength of the component of the weight matrix associated with task *µ*, could enable selective activation of a desired task’s dynamics. Such a modulation can be interpreted as increasing the effective recurrent strength of one task-specific latent dynamical system. Targeted gain control in which inputs lead to changes in effective connectivity through a nonlinearity, neuromodulatory signals, or short-term synaptic plasticity are potential biological substrates (see Discussion).

We develop an analytical theory that characterizes the dynamical regimes of multi-task networks with large *P* and predicts when task selection due to modulation of *D*^(*µ*)^ is successful. The theory applies to networks for which *P* and *N* are large, but with a fixed ratio *α* = *P*/*N*. It also assumes that *R*, the dimension of the manifold associated with a single task, is fixed. This scaling is motivated by the hypothesis that neural circuits perform many distinct tasks, each of which is associated with a low-dimensional manifold (Fig. 3A).

**Figure 3:**
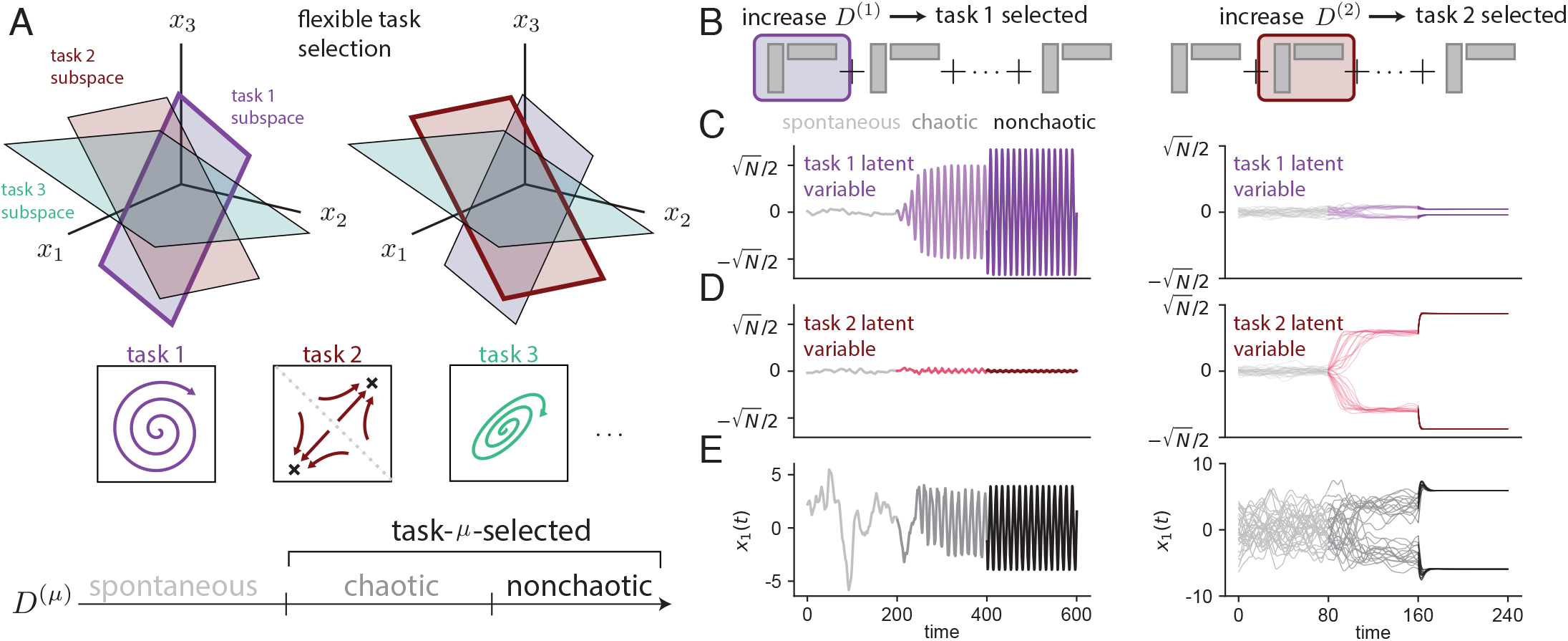
Task-selected states. **(A)** Top: Illustration of task selection as promoting activity in the chosen task’s subspace, which contains task-specific dynamics. Bottom: Phase diagram as a function of the task strength *D*^(*µ*)^, comprising the spontaneous, chaotic task-selected, and nonchaotic task-selected states. Schematic of task selection at the level of connectivity: modulating the strength of the chosen task’s connectivity component through the associated *D*^(*µ*)^ value. **(C)** Trajectory of a task-1 latent variable (normalized by *D*^(*µ*)^), 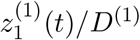, when task-1 limit-cycle dynamics are selected (left) vs. when task-2 bistable dynamics are selected (right), in all 3 regimes. To illustrate bistability, multiple trajectories with different initial conditions are plotted on the right. **(D)** Same as (C) but for a task-2 latent variable 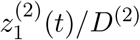. **(E)** Same as (C) and (D) but for an example neuron’s activity.

The theory reveals that modulation of *D*^(*µ*)^ indeed supports selection of task-related dynamics in multi-task networks defined as Eq. 4. These connectivity modulations can place the dynamics of such networks into one of four distinct regimes. In the first, weights are small in magnitude and no activity is produced, so we do not consider this case further. In the second, the *spontaneous* state, chaotic fluctuations occur and no task is dominant (as in Fig. 2E). The other two states are the chaotic and nonchaotic *task-selected* states. The network transitions from the spontaneous state to a state in which task *µ* is selected if *D*^(*µ*)^ is increased sufficiently (Fig. 3B). Of course, task selection could be trivially achieved by setting *D*^(*µ*)^ to zero for all but one task, or by increasing it for a single task so dramatically that it dominates the entire weight matrix. However, our model requires only small increases in *D*^(*µ*)^ to produce large changes in task-related dynamics (see Discussion).

The simulations thus far have depicted the transition from a task-selected state to the spontaneous state as *α* is varied (Fig. 2D). Henceforth, we consider transitions from the spontaneous state to task-selected states due to modulation of *D*^(*µ*)^ for fixed *α*. Such a transition can be induced for any task by modulating its associated *D*^(*µ*)^ value (assuming the task does not involve purely decaying dynamics; see Methods). Thus, there are many distinct task-selected states, for which activity is large in the selected task’s subspace and small in the others (Fig. 3C,D). This contrasts with networks without modulation of *D*^(*µ*)^, in which winner-take-all dynamics do not permit the selection of different tasks (Fig. 2).

### Theory of the spontaneous state

The spontaneous state occurs when two conditions on the connectivity are met. First, for the network not to be quiescent, the average synaptic weight magnitude, 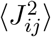, must be sufficiently large (see Methods). This is similar to the condition for spontaneous activity to occur in networks with independent and identically distributed weights [36] and requires that *α* > 0, i.e. the number of tasks indeed scales with the network size. The second condition is that there must be no task *µ* whose strength *D*^(*µ*)^ is so large that it dominates the other tasks, leading to a task-selected state.

Here we describe latent-variable dynamics, and in later sections, we describe single-neuron activity (the theory describes both; see Methods). In the spontaneous state, the latent variables 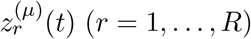 for each task (*µ* = 1, …, *P*) follow

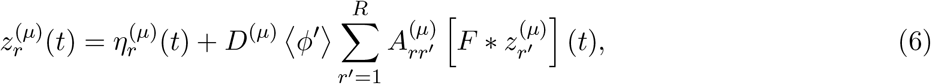

where 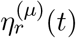 is a temporally correlated Gaussian noise process and *F*(*τ*) is a temporal filter. Eq. (6) describes the dynamics in previous simulations when no task is dominant (e.g. Fig. 2E; or Fig. 2G, bottom).

Eq. (6) can be compared to Eq. (3), which describes a single-task network (Fig. 2A). The first difference is the presence of an additional term 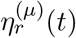, which represents interference due to effective noise from the many small overlaps with other tasks’ dynamics. The second is that the gain factor ⟨*ϕ*′⟩ is time-independent. This reflects the fact that, in the spontaneous state, the gain is not entrained to any single task’s latent variables. Instead, it is determined by an average over many vanishingly correlated latent variables of different tasks, producing a time-independent quantity (Fig. 2G, black vs. gray). Finally, the linear filter *F*(*τ*) is in general different from that of Eq. (3); it depends on the statistics of the ensemble of *D*^(*µ*)^ and **A**^(*µ*)^ (see Methods).

Therefore, in the spontaneous state, the dynamics of 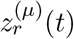, for any task *µ*, are noise-driven and linear. As such, nonlinear dynamics such as asymptotically stable limit cycles or multistability cannot occur. Nevertheless, the spontaneous-state dynamics within a single task subspace are related to those that would be produced if the task were dominant (Fig. 2A,B). For example, the latent variables associated with limit-cycle dynamics exhibit an oscillatory component (Fig. S5A). Similarly, the latent variables associated with bistability exhibit a slow activity mode along the same axis that contains the fixed points in a single-task network (Fig. S5A). Thus, in the spontaneous state, latent variables for all tasks simultaneously follow noisy, linearized, and weakened versions of their task-selected dynamics.

### Theory of the task-selected state

When a specific task *µ*’s strength *D*^(*µ*)^ is increased sufficiently, the linear dynamics of the spontaneous state (Eq. 6) become unstable, leading to rapid growth of the task’s latent variables. When this occurs, the gain factor ⟨*ϕ*′(*t*)⟩ is no longer time-independent and instead is dynamically entrained to task *µ*’s latent variables. This change amounts to the selection of the nonlinear dynamics associated with these variables and the suppression of unselected task dynamics (Fig. 3A). Just above this value of *D*^(*µ*)^, task-selected dynamics coexist with chaotic fluctuations. Increasing *D*^(*µ*)^ further leads to a second transition, beyond which the dominant task completely suppresses chaotic fluctuations, creating a nonchaotic task-selected state. Previous studies have identified related transitions in other network models [28, 37–40].

How does the activity of single neurons and of task-related latent variables change across these transitions? Increasing *D*^(1)^ or *D*^(2)^ (Fig. 3B) beyond the first transition threshold—to place the network in the corresponding chaotic task-selected state—leads to the associated latent variables growing in magnitude (from 𝒪 (1) in the spontaneous state to 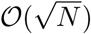, while latent variables for any non-selected task remain 𝒪 (1) (Fig. 3C,D). In the chaotic task-selected state just above the transition, the activity of each neuron exhibits a combination of tuning to the selected task’s latent variables and chaotic fluctuations, with the two effects contributing to single-neuron activity on the same order (Fig. 3E, darker gray). In contrast, the selected task’s latent variables are noiseless to leading order, as single-neuron fluctuations average out in the projection onto the task subspace. Thus, the chaotic task-selected state is characterized by variability at the single-neuron level but not at the level of task-relevant variables. Increasing *D*^(1)^ or *D*^(2)^ further to enter the nonchaotic task-selected state suppresses this variability at the single-neuron level as well (Fig. 3E, black).

The value of *D*^(*µ*)^ required for task selection depends on task *µ*’s structure, captured by its overlap matrix **A**^(*µ*)^, as well as on the number and structure of other tasks (see Methods). Fig. 4 illustrates these dependencies for an analytically tractable class of networks in which each task corresponds to the generation of a stable oscillation with a different frequency (see Methods). A larger value of *D*^(*µ*)^ is required for task selection when the number of tasks is increased (horizontal slices in Fig. 4A; compare with Fig. 2D). The dependence of the critical value on the overlap matrix of the selected task reduces, in this case, to a dependence on the selected task’s oscillation frequency (Fig. 4B).

**Figure 4:**
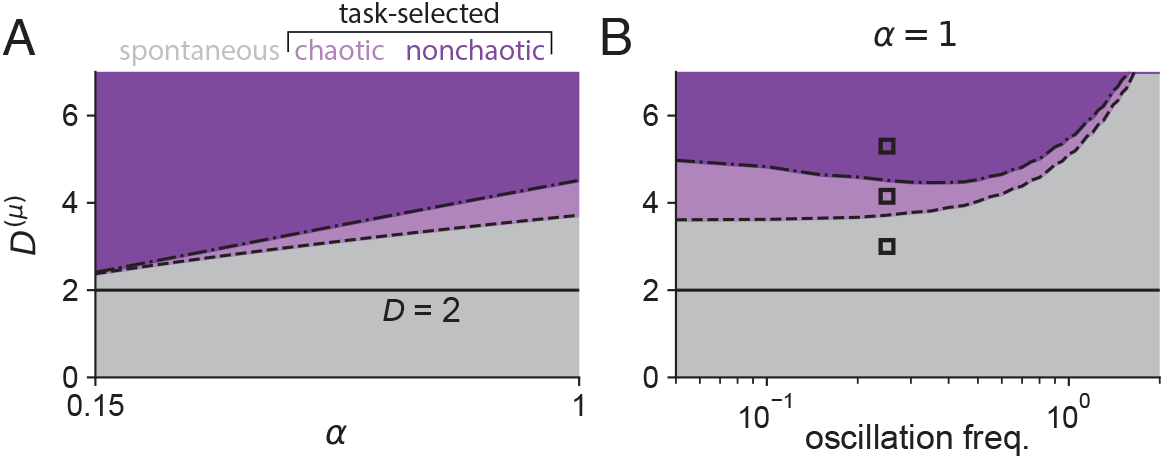
Phase diagrams for networks generated as a sum of single-task components associated with oscillatory dynamics (see Methods). **(A)** Network state as a function of normalized number of tasks *α* = *P*/*N* and the strength of one task *D*^(*µ*)^. All other tasks have *D*^(*ν*≠*µ*)^ = 2 (solid line). Dashed line: Boundary between spontaneous and chaotic task-selected state for task *µ*. Dot-dashed line: Boundary between chaotic and nonchaotic task-selected state for task *µ*. **(B)** Same as (A) but for fixed *α* = 1 and instead varying the frequency of the limit cycle task that is selected. Black boxes: Parameter values whose associated single-neuron properties are examined in Fig. 5.

### Single-neuron statistics in spontaneous and task-selected states

Thus far, we have focused on describing latent variables defined by projections of network activity onto task-specific subspaces. The theory also describes activities of individual neurons (see Methods). In the spontaneous state, the activity of a given neuron *i* is described by

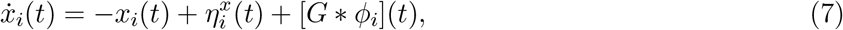

where 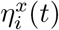 is a temporally correlated Gaussian noise process and *G*(*τ*) is a filter that produces a nonlinear self-coupling of each neuron to its firing rate. Each neuron’s activity fluctuates with negligible tuning to any single task’s latent variables (Fig. S5A). The statistics of these fluctuations are homogeneous across neurons. The theory allows analytic calculation of the correlation functions of 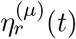 and *η*^*x*^(*t*) and the kernels *F*(*τ*) and *G*(*τ*). This leads to predictions for the single-task and single-neuron correlation functions, 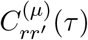 and *C*^*ϕ*^(*τ*), respectively, which we verify with simulations (Fig. S5).

In task-selected states, the equation describing single-neuron activity is similar to Eq. (7), with an additional term corresponding to the neuron’s tuning to the dominant task’s latent variables:

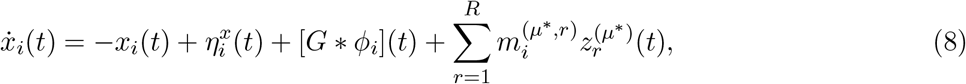

where *µ*^*∗*^ is the index of the dominant task. The 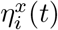 term is still a Gaussian noise process, but its kernel is not time-translation-invariant and instead depends on the dynamics of the dominant task (see Methods).

In task-selected states, neurons are statistically heterogeneous, following different trajectories depending on their tuning to the dominant task’s latent variables (Fig. 5A). This heterogeneity implies that the extent to which single-neuron activity can be explained by one task’s latent variables varies across neurons in the population. The mean of this distribution is low in the spontaneous state, higher in the chaotic task-selected state, and highest in the nonchaotic task-selected state (Fig. 5B).

**Figure 5:**
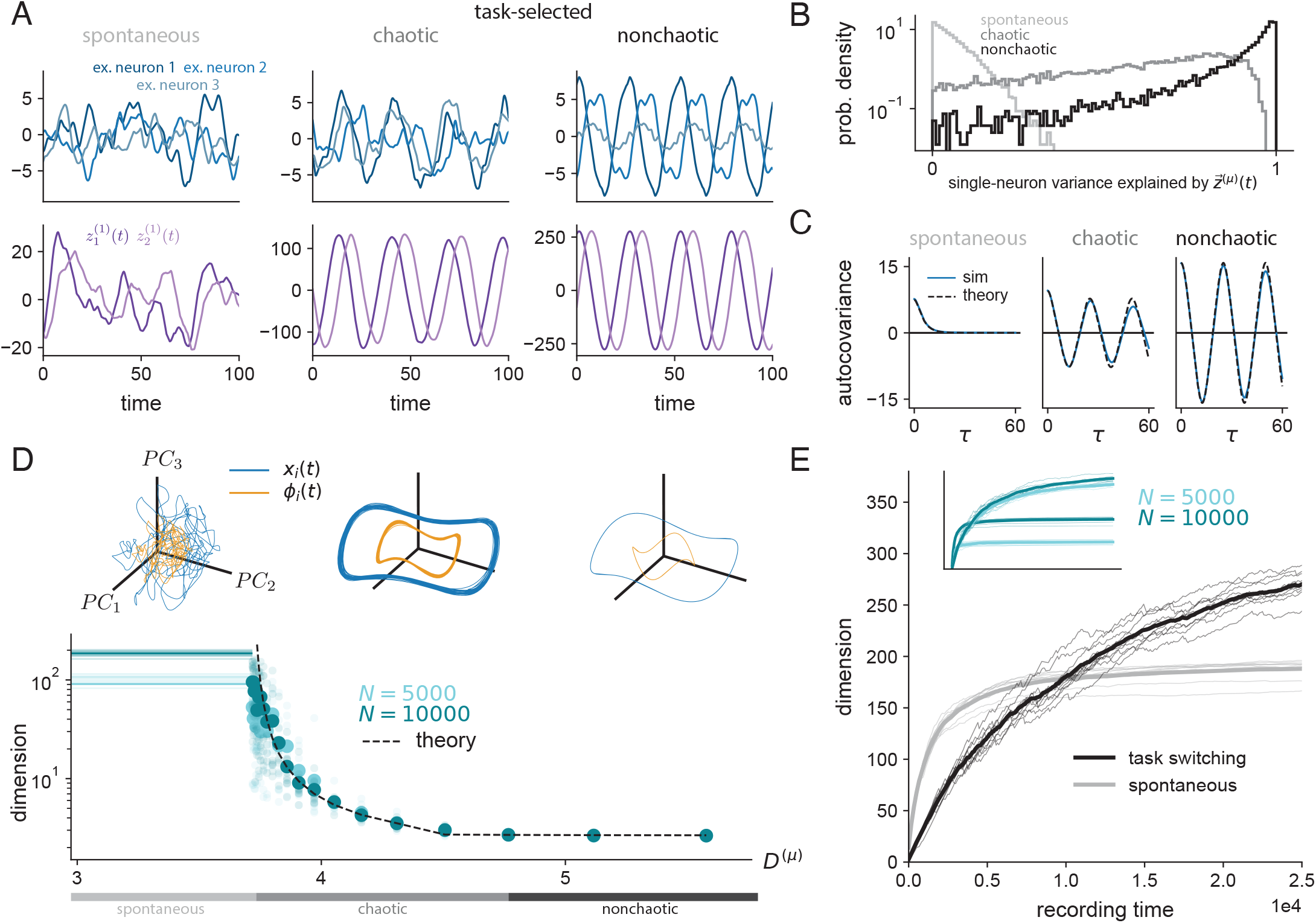
Single-neuron and population properties of a multi-task networks. Parameters are the same as Fig. 4 (see Methods). **(A)** Example neurons (top) and latent variables for the selected task (bottom) across network states. **(B)** Histogram *R*^2^ values for linear regression of *x*_*i*_(*t*) against task-*µ* latent-variable activity. **(C)** Average single-neuron autocovariance functions *C*^*x*^(*τ*) = ⟨*x*_*i*_(*t*)*x*_*i*_(*t* + *τ*) ⟩ _*t,i*_. **(D)** Top: Principal component trajectories across same network states from (A), (B), and (C). For currents *x*_*i*_(*t*) and firing rates *ϕ*_*i*_(*t*), principal component axes and projections onto these axes are computed separately, with *ϕ*_*i*_(*t*) projections doubled for visibility. Axis scales: 150 (left), 300 (middle), 500 (right). Bottom: Linear embedding dimension computed as participation ratio of the firing rates *ϕ*_*i*_(*t*), as a function of *D*^(*µ*)^. Light dots: individual network simulations. Heavy dots: averages over networks. **(E)** Firing-rate dimension as a function of recording time for a network in the spontaneous state and switching across task-selected states. For the latter, one of 150 tasks is randomly selected for 80 time constants, followed by 3 time constants of spontaneous activity, and this process is repeated. Thin lines: individual network simulations. Thick lines: average. Inset: The same plot but for networks with different *N*, showing dependence on *N* in the spontaneous state.

The average autocovariance function *C*^*x*^(*τ*) = ⟨*x*_*i*_(*t*)*x*_*i*_(*t* + *τ*)⟩_*t,i*_ illustrates the differences between the temporal statistics of single-neuron firing rates in the spontaneous and task-selected states (Fig. 5C). The first transition occurs when task-related structure, in this case oscillations, becomes visible at the single-neuron level (Fig. 5C, left vs. middle). The second transition occurs when all firing-rate variance is accounted for by dominant-task activity, or equivalently, in the case of a limit-cycle task, when *C*^*x*^(0) = *C*^*x*^(*T*) for an oscillation with period *T* (Fig. 5C, middle vs. right).

### Dimensionality of spontaneous and task-selected states

To analyze the structure of population activity in multi-task networks, we also calculate its dimension. We quantify dimension using the participation ratio, a measure of linear dimensionality that approximates the number of orthogonal directions in activity space that are explored by the network’s dynamics. This quantity has been the subject of numerous experimental and theoretical studies [37, 41–45].

In the spontaneous state, the dimension scales with the network size *N*, as previously observed in networks with random connectivity (43; Fig. 5D, left). Although this scaling produces dimension values that are large, they are always much lower than the ambient dimension *N* due to weak correlations among neurons [45], a common observation in neural data [22, 46, 47]. At the other extreme, in the nonchaotic task-selected state, the dimension is determined solely by the dominant task’s dynamics and is therefore proportional to *R* (Fig. 5D, right). Between these extremes, in the chaotic task-selected state, the dimension is larger than this limiting value by a factor that does not grow with *N* and can be predicted by the theory (Fig. 5D, middle; see Methods). This increase arises from chaotic fluctuations in directions orthogonal to the dominant task’s subspace. The magnitudes of these fluctuations depend on the values of *D*^(*µ*)^ for both the selected and unselected tasks (Fig. S6).

These results suggest that experimental measurements of neural dimensionality should be interpreted in the context of the underlying network state. In particular, the observed dimension may depend on which task-selected states are present, the statistics of transitions between these states, and the dimension of the manifolds associated with each individual task. Consistent with previous studies of neural activity during specific behavioral tasks [21, 48, 49], our model predicts that the dimension of activity is low when activity is measured within a single task-selected state, even when fluctuations are present at the single-neuron level. An experimentally observed increase beyond this value can arise through two distinct mechanisms. First, the network may transition into the spontaneous state, whose dimension scales with network size, consistent with analyses of large-scale neural recordings [20]. Second, the dimension of activity may increase through the sequential engagement of multiple task-selected states, each associated with a different low-dimensional manifold.

These mechanisms make distinct predictions for how the dimensionality of activity grows with recording time (Fig. 5E). In the spontaneous state, dimensionality increases rapidly at short times and then saturates (Fig. 5E, gray). When the network sequentially switches across task-selected states, the rate of dimensionality growth depends on the amount of time spent in each task state. Even in cases in which this time is short (as in our simulations), dimensionality grows more slowly than in the spontaneous state (Fig. 5E, black).

The two mechanisms also differ in their saturation values. In the spontaneous state, network activity explores a chaotic attractor in which projections onto certain directions in neural activity space contain more variance than others, limiting dimensionality. In contrast, sequential task selection produces a sequence of trajectories that are highly decorrelated from one another. As a result, the saturation value for sequential task selection is approximately equal to the number of visited task states multiplied by the within-task dimension. This value can be either higher or lower than the spontaneous-state dimension. For a sufficiently large number of tasks, the dimensionality associated with sequential task selection can greatly exceed that of the spontaneous state (Fig. 5E, inset). Altogether, the theory demonstrates that either a fixed high-dimensional covariance matrix or state-dependent switching among different low-dimensional covariance matrices are both potential models for high-dimensional activity in neural recordings.

## Discussion

We identified limitations to multi-task capabilities in nonlinear low-rank networks due to interference among tasks. Such limitations are not evident in analyses of networks constructed to perform a single task or in linear networks that lack the winner-take-all interference we described. We proposed a solution to the problem of interference that relies on modulation of effective connectivity. Networks with such a mechanism transition between a spontaneous state in which no task is selected and task-selected states in which activity lies primarily on a task-specific low-dimensional manifold. Our theory thus retains the analytic tractability of low-rank networks [28, 29, 34] while describing the geometry of neural dynamics associated with multiple tasks.

### Neuronal architectures for multi-task networks

The connectivity of our model, and its modulation during task selection, may be interpreted in multiple ways. If the network is viewed as a single recurrently connected circuit, modulation of *D*^(*µ*)^ corresponds to a change in the strength of a single task-related component of the weight matrix (Fig. 6A). Such a change requires selective modulation of synapses depending on their participation in this weight matrix component. Potential biological substrates include rapid synaptic plasticity or targeted neuromodulation [51–56].

**Figure 6:**
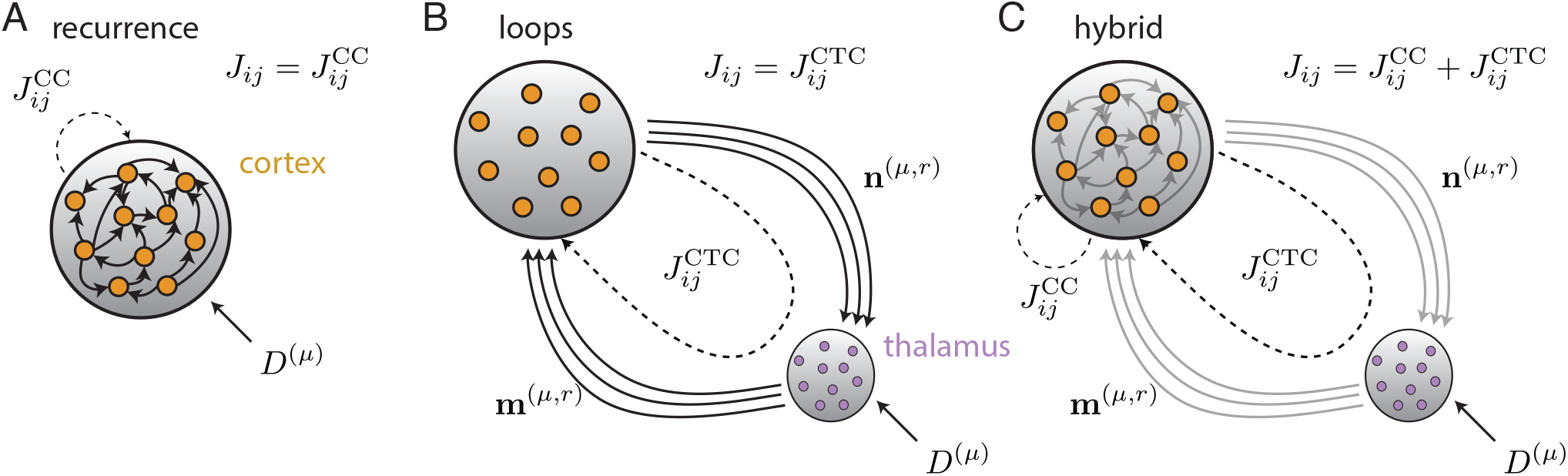
Illustrations of three interpretations of the network’s effective connectivity. **(A)** Modulation of *D*^(*µ*)^ modifies the recurrent connections 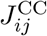 of a cortical network. **(B)** Interactions between neurons are driven by effective connectivity 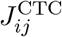 defined by thalamocortical loops. Modulation of *D*^(*µ*)^ adjusts the gain of thalamic neurons [50]. **(C)** Model in which the effective connectivity is a combination of (A) and (B).

Alternatively, the loadings **m**^(*µ,r*)^ and **n**^(*µ,r*)^ may be interpreted as synaptic weights between a network and an auxiliary population of neurons, for instance thalamocortical and corticothalamic weights, respectively (Fig. 6B). In this case, task selection is accomplished through adjustment of the gain of a subset of task-associated thalamic neurons. A linear network model of such a mechanism has been proposed [50]. More generally, because only small modulations of effective connectivity are required in our nonlinear model, its weights may be interpreted as a combination of fixed corticocortical and adjustable thalamocortical synapses (Fig. 6C).

We focused on how a single network can generate many distinct sets of low-dimensional dynamics, without explicitly modeling task-related sensory inputs or how these dynamics are transformed to produce behavior. In low-rank single-task networks, task-related inputs define additional dimensions of the task subspace [28, 34]. Similarly, we expect our qualitative results on interference and winner-take-all dynamics to generalize to cases where distinct tasks operate with distinct input and output channels. Future work could examine the consequences of sharing neural modes across tasks, whether input channels, latent dynamical modes, or output channels [2, 3, 5, 7].

### Related models of multi-tasking

Previous studies have examined multi-tasking in recurrent neural networks that are trained end-to-end to perform individual tasks that depend on a contextual input [3, 27], or sets of tasks commonly used in neuroscience experiments [4, 14]. These models produce diverse dynamical behaviors but their weight matrices have complex statistical structure from the training process that limits analytical tractability. It would be interesting to examine whether the tonic inputs commonly used to signal task context in trained multi-task networks effectively drive similar task-selection mechanisms via gain modulation, perhaps in task-specific populations [14, 29, 34], by shifting the operating point of the nonlinearity.

Other works have studied nonlinear models with closed-form expressions for the weight matrix, including summations over a large number of low-rank components, as in our model. In certain cases, such connectivity leads to chaotic activity [38, 45, 55, 57, 58]. These studies considered networks that are designed to produce specific dynamical features, including fixed points [38], continuous attractors [58], and sequential activity [55, 57]. Our model, on the other hand, permits each task to be associated with any low-dimensional dynamical system described by Eq. (3), that is, any dynamical system that can be described within the framework of single-population low-rank networks [29]. Thus, although we limited our analysis to limit cycles and fixed points, our model also accommodates sequential activity and non-normal latent dynamics [59].

Modulating connectivity has been proposed in previous works. A previous study developed a linear network model of thalamocortical interactions (Fig. 6B) that sequences transient motor motifs [50]. Other models have used statistical inference to learn time-varying scaling factors that adjust low-rank connectivity components in nonlinear networks [6, 56]. Our theory provides an analytical account of such modulation and its consequences for task selection and spontaneous activity.

In biological and artificial systems, complex behaviors are often built by combining simpler behavioral components [14, 60]. In our model, each “task” should be interpreted as a minimal computational component associated with a behavior, which may only be part of what is often referred to as a task in behavioral experiments. For example, fixation, stimulus, delay, and response periods may each demand different network dynamics [6]. To retain analytic tractability, here we assumed no overlap or reuse of dynamics across tasks, but including such structure is an interesting direction for future work.

Related theoretical work suggests that selecting among low-rank connectivity components, each suited to a particular subtask, is a more effective strategy for continual learning than treating each compound behavior as its own learning problem [6]. Animal experiments indicate that complex motor control arises from sequencing different low-dimensional dynamics, each associated with a particular manifold of state space, to generate flexible behavioral sequences [2]. Our theory provides an analytically tractable mechanism for such sequential composition. In contrast, the winner-take-all dynamics of our model does not support multiple simultaneously active tasks, due to the Gaussian-distributed loadings. Such dynamics require additional circuit structure, such as multiple neuronal populations (e.g. from mixture-of-Gaussians loadings) with population-specific gain factors [29, 34]. Nonetheless, the same gain-dependent mechanisms that enable task selection and suppression in the Gaussian case may also facilitate simultaneous selection of multiple tasks—and suppression of all unselected tasks—in more complex models, which we leave to future work. Such models would additionally allow for more complex single-task dynamics beyond what can be achieved with one population [34].

### Magnitude of effective connectivity modulation

Our theory shows that, in nonlinear networks, a modest modulation of effective connectivity is often sufficient for task selection. Adjustment of all values of *D*^(*µ*)^, such that each value is either zero for unselected tasks, or nonzero for the selected task, would achieve task selection trivially by completely overwriting the synaptic weight matrix each time a task is selected. In contrast, our model requires adjusting the *D*^(*µ*)^ value for only the selected task, which corresponds to changes in synaptic weights that are 𝒪 (1/*P*), whereas each synaptic weight is 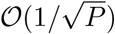. Thus a small task-related connectivity component can dramatically influence a network’s activity, an observation that has been made in models with both structured low-rank and unstructured weight matrix components [28, 33, 39, 61]. Our model demonstrates that this feature holds in networks for which all connectivity components are task-related.

Moreover, the degree of modulation, even relative to 1/*P*, can be made small in our model. The change in *D*^(*µ*)^ required for task selection depends on its distance from the critical value at which task selection occurs (Fig. 4B, black solid line vs. dashed). When *α* is small, a plausible limit in which the number of tasks is large but much smaller than the number of neurons, these distances can be decreased (Fig. 4A). Rapid reconfiguration of dynamics thus occurs with only small connectivity adjustments, as each task is close to selection. In such networks, dynamics are near the edge of stability in the spontaneous state [62], providing a functional explanation for the presence of irregular spontaneous activity.

### Dimensionality and the relationship between spontaneous and evoked activity

Spontaneous activity has historically been defined as activity in the absence of sensory stimulation [63]. Classic theoretical work showed how such activity could be self-generated by a nonlinear network without external inputs [36]. However, recent high-yield recordings during periods of unstimulated, uninstructed behavior have revealed high-dimensional activity whose high-variance components are tuned to kinematic variables [19, 20], which is not well described by unstructured chaotic activity. The spontaneous state of our model has similarities to the chaotic regime of unstructured networks [36] but describes both the irregular activity of neurons and also of task-relevant latent variables. The spontaneous state of our model may explain the idiosyncratic behaviors that have been observed to accompany periods of “quiet wakefulness” [15], while the chaotic task-selected state may explain how such behaviors coexist with coherent goal-directed actions [64].

When freely exploring, rodents perform a variety of self-initiated behaviors, such as whisking or grooming [1]. Many of these behavioral “syllables” feature precise kinematic control that is likely not explained by the spontaneous state of our model. Rather, recordings during long periods of free behavior may correspond to switching among different task-selected states of our model. This predicts that individual neurons should have heterogeneous tuning to kinematic variables for an active behavioral syllable, and this tuning will change across syllables (Fig. 5B). A second prediction is that, when pooling neural activity across distinct syllables, the observed dimension should increase with the diversity of behavioral syllables observed. Conversely, dimensionality measured over many repetitions of a single behavioral syllable should be small, even if there is trial-to-trial variability at the single-neuron level (Fig. 5D). A final prediction is that the initial rate of dimension growth relative to recording time should be faster during quiet wakefulness than during behavior, and the magnitude of this effect depends on the average syllable duration (Fig. 5E). These predictions can be examined in neural recordings during free behavior [65].

## Acknowledgments

We thank Manuel Beiran for discussions and for proposing the inhibition-stabilized model of Fig. S3. We thank Lea Duncker for feedback on the manuscript. OM, DGC, and ALK were supported by the Kavli Foundation and the Gatsby Charitable Foundation (grant no. GAT3708). ALK was also supported by the National Science Foundation (grant no. 2443158).

## Methods and Materials

### Dynamical mean-field theory

We now describe the full theory in greater detail, leaving derivations of the results to the Appendix. The dynamics describing the recurrent network (Eq. 1) can be interpreted as a bidirectionally coupled system of neurons and latent variables (Fig. 7A). Each latent variable is defined as a sum over neuronal firing rates, and in turn the input to each neuron can be written as a sum over latent variables:

**Figure 7:**
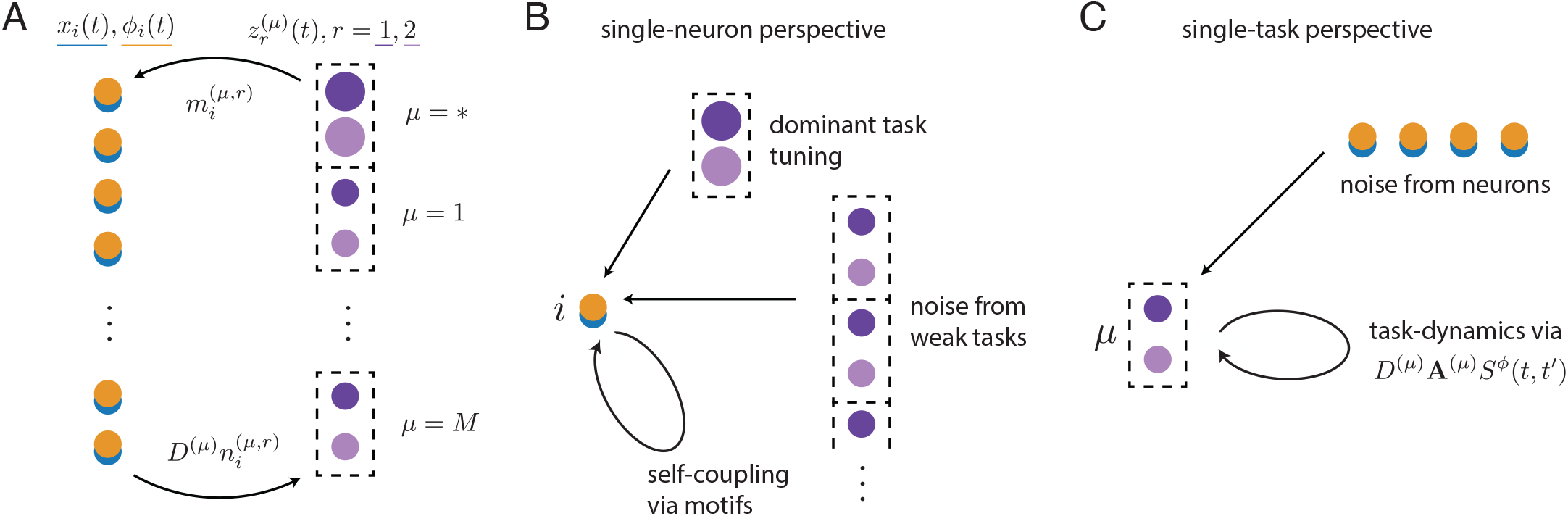
Conceptual illustration of the DMFT. **(A)** Cartoon illustrating an alternative interpretation of the neuron-to-neuron recurrent dynamics, as continuous-time, bidirectional interactions between a reservoir of neurons (blue: currents; yellow: firing rates) and a reservoir of latent variables (dark and light purple). Here we have a single set of dominant latent variables (*P*^*∗*^ = 1), illustrated as larger purple disks. (**B)** Types of effective input a single neuron receives in the DMFT: input from dominant task latent variables with tuning determined by 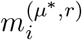 (top); summed input from weak-task latent variables, which is well described as a sample 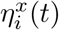 from a Gaussian process (middle); and nonlinear effective self-interaction that arises through coherent summation of the neuron’s perturbative effect on latent-variable trajectories (bottom). (**C)** Types of effective input a single weak-task latent variable receives in the DMFT: summed input from neurons, which is well described as a sample 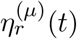 from a Gaussian process (top); and effective self-interactions that are mediated by single-neuron response properties as well as task-specific parameters *D*^(*µ*)^ and **A**^(*µ*)^ (bottom).

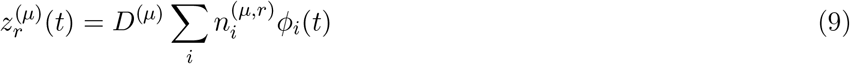

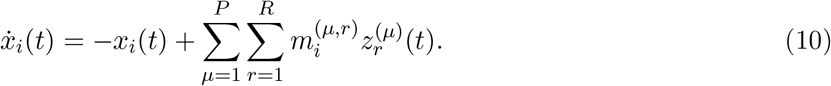

The magnitudes of these latent variables can be large 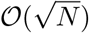, as in Fig. 2A) or small (𝒪 (1), as in Fig. 2E). We refer to any task whose latent variables are large as a “dominant” task, and the others as “non-dominant” tasks. This distinction is central to the mean-field description, because each neuron’s activity is significantly affected by any dominant task, meaning that a finite fraction of its variance can be explained by that task’s variables. This is not true for the non-dominant tasks, although they collectively provide non-negligible input as an ensemble. A task can be dominant for two reasons: 1) initial conditions or inputs pushing the state of the network to preferentially align with the task subspace, or 2) self-amplifying dynamics within the task subspace, as shown in Fig. 3. A random network state results in weak latent variables for every task.

We set aside a separate index *µ*^*∗*^ = 1^*∗*^, …, *P*^*∗*^, with *P*^*∗*^ ≥ 0 small, to enumerate tasks 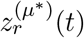 with dominant 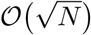 activity, if there are any.^1^ Then *µ* = 1, …, *P* continues to enumerate the many non-dominant tasks with 𝒪 (1) activity. In the *N, P* → ∞ limit with fixed ratio *α* = *P*/*N*, fixed *P*^*∗*^, and fixed *R*, we can statistically describe the trajectories of all neurons and weak-task-related latent variables as decoupled processes, rather than as solutions to the high-dimensional dynamics of Eq. (1). In this mean-field description, the activity of the *i*th neuron *x*_*i*_(*t*) and the *r*th latent variable 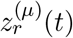 of the *µ*th non-dominant task evolve as

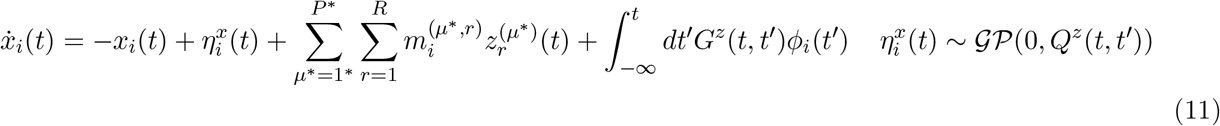

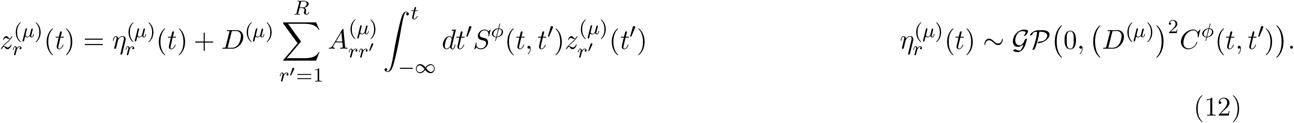

In deriving the theory, several functions of time emerge that are shared by all neurons and tasks: four kernel functions *C*^*ϕ*^(*t, t*′), *S*^*ϕ*^(*t, t*′), *Q*^*z*^(*t, t*′), *G*^*z*^(*t, t*′) that describe effective noise statistics and interactions, plus the set of dominant-task latent variable trajectories 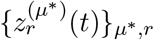. (In the main text, we wrote *S*^*ϕ*^(*t, t*′) = ⟨*ϕ*′(*t*)⟩ *F*(*t, t*′) and *G*^*z*^(*t, t*′) = *G*(*t, t*′) to simplify notation—see section on self-consistency conditions.) All these functions’ values depend collectively on 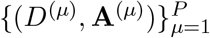 and *α*, and they can be calculated by enforcing self-consistency, as we describe later.

Each neuron *i* is described in Eq. (11) as leaky integration of 3 current sources (Fig. 7B):

1. A Gaussian process noise term 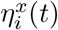 arising from interactions with the large reservoir of weak-task-related latent variables and capturing the effects of chaotic fluctuations. Its temporal statistics *Q*^*z*^(*t, t*′) are inherited from average statistics of non-dominant-task latent variables.
2. Drive from the dominant-task latent variables 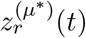 via the neuron-specific loadings 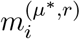.
3. A nonlinear self-coupling to the neuron’s *own* firing rate, which is generally a consequence of introducing task-related overlaps. The kernel mediating this coupling *G*^*z*^(*t, t*′) is determined by weighted averages of the induced network connectivity motifs of all orders.

Similarly, each weakly active task *µ* is described in Eq. (12) as an integral equation with two input sources (Fig. 7C):

1. A Gaussian process noise term 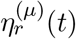 that arises from the sum over many irregular neuronal activity trajectories via Eq. (9). Its temporal statistics (*D*^(*µ*)^)^2^*C*^*ϕ*^(*t, t*′) are determined by the average neuronal autocovariance function and scaled by *D*^(*µ*)^.
2. A coupling among task *µ*’s latent variables mediated by *D*^(*µ*)^**A**^(*µ*)^ and the kernel *S*^*ϕ*^(*t, t*′). The kernel *S*^*ϕ*^(*t, t*′) is determined by average impulse-response properties of the neuronal population. The average gain is a crucial factor in determining *S*^*ϕ*^(*t, t*′), since (un)saturated neurons are less (more) responsive to inputs.

The dominant-task latent variables satisfy

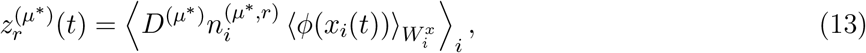

where 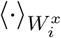 indicates averaging over the random process described by Eq. (11) for a given neuron *i*. In general, this expression does not have a simple closed-form solution, and calculating the forward dynamics requires iterative numerical approaches to maintain self-consistency. However, in special cases where currents *x*_*i*_(*t*) are Gaussian, the dynamics of 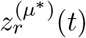 essentially reduce to the simple form of Eq. (3) from the single-task network case, by Stein’s Lemma.^2^

In summary, our theory is able to completely describe network activity for an arbitrary *α* and ensemble of task dynamics, at the level of both single neurons and task-specific subspaces. Inputs to each can be decomposed into regular, task-related signals and effectively noisy fluctuations due to chaos, with mutually determined statistics that can be analytically solved for.

### Note on *N*-scaling

Here we summarize how various structural and dynamical quantities scale with the network size *N*. Components of loading vectors (**m**^(*µ,r*)^, **n**^(*µ,r*)^ for *r* = 1, …, *R, µ* = 1, …, *P*) are 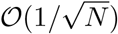, resulting in elements of each single-task weight matrix (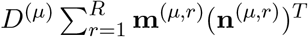 for *µ* = 1, …, *P*) being 𝒪 (1/*N*). Because the number of tasks *P* is 𝒪 (*N*), each single-task weight matrix has zero mean, and the single-task weight matrices are sampled independently, the elements of the full weight matrix that sums over all single-task matrices is 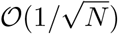.

Neural activities (*x*_*i*_(*t*), *ϕ*_*i*_(*t*) for i = 1, …, N) are 𝒪 (1). Projections of neural activity onto single-task subspaces 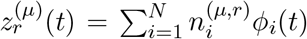 for tasks *µ* = 1, …, *P* and within-task components *r* = 1, …, *R*) scale in two possible ways depending on whether a task *µ* is dominant or non-dominant. In particular, projections onto non-dominant task subspaces are 𝒪 (1), while projections onto dominant-task subspaces are 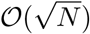. This is because, in the dominant-task case, the network state preferentially aligns with their associated subspaces, and the overall state norm ∥***ϕ***(*t*)∥ is 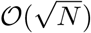 while the norms of loadings vectors are approximately 1. In the language of associative-memory networks that store patterns as fixed points, these forms of (non-)alignment are referred to as uncondensed or condensed.

These distinctions in scaling are visible when comparing networks of different sizes (Fig. 2D light vs. dark cyan). The activity level of dominant task *µ* = 1 is scaled by approximately 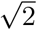 when the number of neurons is doubled, but past the transition to chaos—when task 1 loses dominance—its activity level drops to a value that is the same for networks of different sizes.

### Connectivity statistics

The free parameters of the model are *P, R*, and 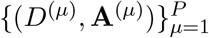. Statistics of *J*_*ij*_ can be written in terms of these parameters. The overall strength of the connectivity, which we call *g*_*eff*_, is given by the variance of the off-diagonal elements of **J** and depends on the second moment of the task weights (but not the overlaps):

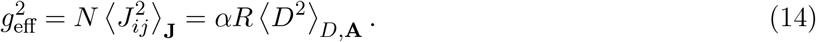

Like the analogous *g* parameter of [36], holding other parameters fixed while increasing this degree of “disorder” eventually effects a transition to chaotic activity, with stronger chaotic fluctuations with increasing *g*_*eff*_. However, in this model with connection symmetries, *g*_*eff*_ does not alone determine the network dynamics.

The mean connection strength is

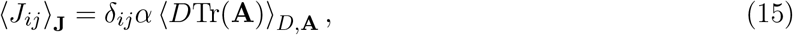

implying strong autapses *J*_*ii*_ ∼ 𝒪 (1) when the overlaps have a nonzero average trace. This homogeneous mean autapse strength ⟨*J*_*ii*_⟩_**J**_ can be interpreted as a zeroth-order connectivity motif *ρ*_0_. The first-order connectivity motif strength *ρ*_1_ is the correlation between transposed elements for *i*≠ *j*,

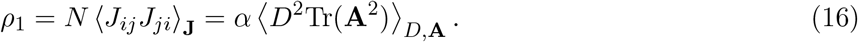

Higher-order connectivity motifs, e.g. *ρ*_2_ = ⟨*J* _*ij*_ *J*_*jk*_ *J* _*ki*_ ⟩_**J**_, arise from nonzero average traces of higher matrix powers of **A**.

If present, these symmetries will lead to effective self-coupling between each neuron and itself, whether through strong self-connections or multi-synaptic pathways. In a Neumann expansion of the the spontaneous state self-coupling kernel *G*^*z*^(*ω*) (see section on self-consistency conditions),

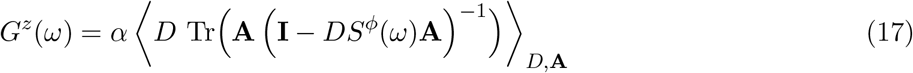

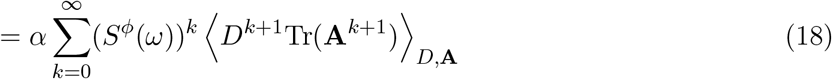

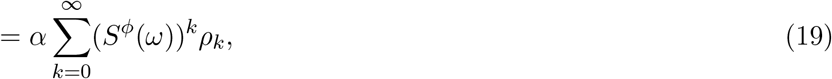

which matches our intuition that neurons self-interact through strong network motifs, modulated by the appropriate power of the impulse-response function.

This self-coupling is responsible for non-Gaussian currents *x*(*t*), which makes calculations difficult. However, designing a non-trivial ensemble of dynamics that eliminates *ρ*_*k*_ for every *k* is possible, and in such cases we get Gaussian *x*(*t*).

### Stability condition for 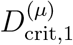

Computing the stability condition for an integral equation of the form

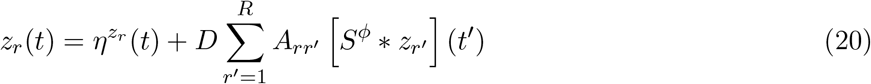

requires the Laplace transform of the transfer function **M**^(*µ*)^(*s*) = (**I** −*D*^(*µ*)^*S*^*ϕ*^(*s*)**A**^(*µ*)^)^*−*1^. All singularities *s*^*∗*^ of this inversion must satisfy ℜ 𝔢 (*s*^*∗*^) < 0 for a value of *D*^(*µ*)^ to produce stable dynamics. This condition is generally difficult to calculate without a closed-form expression for *S*^*ϕ*^(*τ*), as it requires numerical estimation of the Laplace transform. If we assume vanishing connectivity motifs and recover Gaussian currents, implying *S*^*ϕ*^(*τ*) = Θ(*τ*) ⟨*ϕ*′⟩ *e*^*−τ*^, the dynamics become equivalent to a leaky, noise-driven ODE, and the stability condition simplifies to

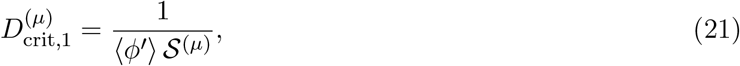

where 𝒮^(*µ*)^ = max[ℜ𝔢 (*λ*) |*λ* eigenvalue of **A**^(*µ*)^] is the spectral abscissa of **A**^(*µ*)^.

### Self-consistency conditions

For the rest of the methods section and appendices, we often drop the neuron index *i* and non-dominant task index *µ*, because in the DMFT they are thought of as ensembles from which *N* and *P* samples are taken, respectively, rather than explicit lists of elements. A given non-dominant task *µ* has an associated set of *D*, **A** values that are sampled from a joint distribution Pr(*D*, **A**), and a given neuron *i* has an associated set of normalized (to 𝒪 (1)) couplings 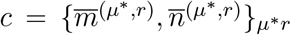 to the dominant-task latent variables. We denote the random process for a single neuron with given couplings c as 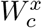, and the random process for a single non-dominant task with given dynamics *D*, **A** as 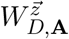. The fully general set of self-consistency conditions are

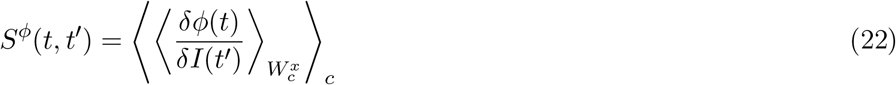

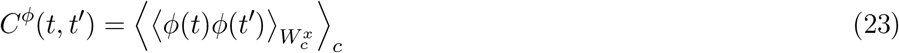

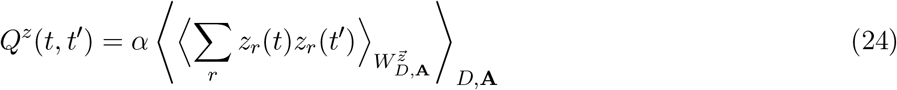

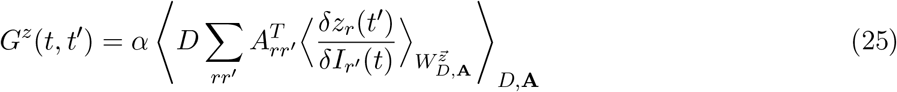

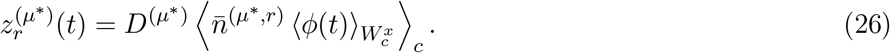

Each kernel function (or order parameter) is computed as a within-process average parameterized by some value, e.g. 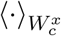 or 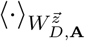, that is itself averaged over the ensemble of all such parameters, e.g. ⟨*·*⟩_*c*_ or ⟨·⟩_*D*,**A**_.

Thus, the DMFT amounts to a bipartite ensemble of neurons and task-related latent variables with mutually referential statistics. The kernels *Q*^*z*^(*t, t*′) and *G*^*z*^(*t, t*′) defining the single-neuron stochastic dynamics are inherited from averages over the ensemble of latent variables, hence their superscripts referencing their definition. Conversely, the kernels *C*^*ϕ*^(*t, t*′) and *S*^*ϕ*^(*t, t*′) defining single-task-latent-variable stochastic dynamics are inherited from averages over the ensemble of neurons. Again, to avoid explaining the bipartite structure implicit in this notation in the main text, we simplified notation and replaced *S*^*ϕ*^(*t, t*′) = ⟨*ϕ*′(*t*)⟩ *F*(*t, t*′) and *G*^*z*^(*t, t*′) = *G*(*t, t*′).

### Spontaneous state self-consistency conditions

In the spontaneous state, the self-consistency conditions simplify tremendously. First, there are no 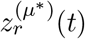 values that need to be solved for, and thus the single-neuron dynamics simplify. Second, we can assume stationarity of the kernel functions, i.e. that each kernel function depends only on *t* − *t*′, not *t* and *t*′ separately. (This is a fairly mild additional assumption, but one that may be broken with sufficiently strong connectivity motifs [66]). Neurons are now statistically homogeneous, so we do not need to average over the ensemble of processes 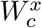 and can instead consider a single representative process 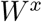. The non-dominant-task latent variables are linear, and their associated autocovariance functions and impulse-response functions can be written compactly in Fourier space. With all of these simplifications, the self-consistency equations take the form

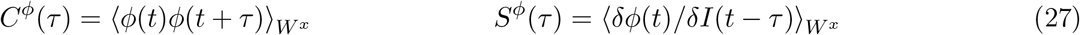

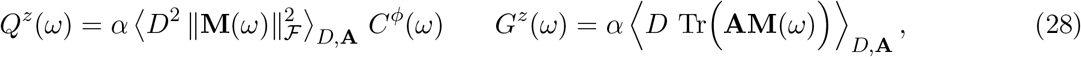

where **M**(*ω*) =(**I** − *DS*^*ϕ*^(*ω*)**A**)^*−*1^ is the transfer function for a task with dynamics according to *D*, **A**.

### Methods for solving the DMFT

To solve the DMFT means to calculate the kernel functions *C*^*ϕ*^(*t, t*′), *S*^*ϕ*^(*t, t*′), *Q*^*z*^(*t, t*′), *G*^*z*^(*t, t*′), plus the set of dominant-task latent variable trajectories 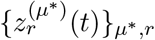 for an ensemble of task dynamics in a given discretization of time.

#### Fully general *t, t*′ case

Solving the theory in this most general case is in principle possible by iteratively enforcing self-consistency in sampling a large number of single-neuron trajectories for provisional guesses of each kernel function and the dominant-task latent variable trajectories. A summary of the process:

1. Start with initial guesses for *C*^*ϕ*^(*t, t*′), *S*^*ϕ*^(*t, t*′), and 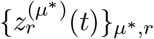
2. From *C*^*ϕ*^(*t, t*′) and *S*^*ϕ*^(*t, t*′), compute *Q*^*z*^(*t, t*′) and *G*^*z*^(*t, t*′) by explicitly averaging over the *P* non-dominant tasks in the ensemble. Since the non-dominant latent dynamics are linear, we can take advantage of having analytic expressions for their within-task cross-covariances and impulse-response functions, which are necessary ingredients for computing *Q*^*z*^(*t, t*′) and *G*^*z*^(*t, t*′).
3. Simulate *N*_*batch*_ samples of Eq. (11) in parallel, each requiring an independent sample of the Gaussian noise *η*^*x*^(*t*) (by the Cholesky decomposition of *Q*^*z*^(*t, t*′)), as well as a sample of the dominant-task coupling strengths 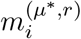.
4. Empirically measure the single-unit autocovariance, impulse-response functions, and dominant-task latent variable trajectories based on these samples. (The autocovariance and dominant latent variables are straightforward to calculate from their definitions, while the impulse-response function requires differentiating the individual *x*_*i*_(*t*) trajectories.)
5. Smoothly update the values of *C*^*ϕ*^(*t, t*′), *S*^*ϕ*^(*t, t*′), and 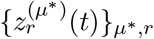 based on these empirical measurements
6. Repeat 2-5 until convergence.

Details for a similar method applied to modern Hopfield networks are outlined in [67].

Although this method, as well as some other methods we present below, is numerical in nature, it still does not require ever constructing or simulating a *network* according to Eq. (1) with a synaptic weight matrix—only sampling neural activity and task-related latent variables as random processes with mutually referential statistics.

We did not ultimately present any results with this method, as it is prohibitively computationally expensive with difficult boundary conditions. However, we summarize it to the reader to emphasize that, as written, the DMFT simply encodes a set of mutually referential statistics between a reservoir of neurons and a reservoir of non-dominant latent variables, with a special set of dominant-task latent variable trajectories. In principle, a self-consistent set of such statistics can be found by iterative sampling and updating. However, calculation becomes much easier in certain special cases, as we articulate below.

#### Spontaneous state case

At a high-level, the general method for solving the theory for the spontaneous state is similar to the fully general method, but now we have much simpler Fourier representations of the kernel functions, rather than explicit *t* and *t*′ dependence, and we do not have to worry about dominant-task latent variable trajectories. The process is simplified to:

1. Begin with an initial guess for *C*^*ϕ*^(*τ*) and *S*^*ϕ*^(*τ*).
2. Based on these values, compute *Q*^*z*^(*ω*) and *G*^*z*^(*ω*) in Fourier space using Eq.(28) with an explicit average over the *D*^(*µ*)^, **A**^(*µ*)^ ensemble.
3. Sample *N*_*batch*_ Gaussian noise samples of 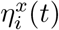 based on the kernel *Q*^*z*^(*τ*), and compute *x*_*i*_(*t*) and *ϕ*_*i*_(*t*) from Eq. (11) without dominant-task latent variables.
4. Empirically measure *C*^*ϕ*^(*τ*) and *S*^*ϕ*^(*τ*), leveraging the Furutsu-Novikov theorem for estimating *S*^*ϕ*^(*τ*): *S*^*ϕ*^(*ω*) = *C*^*ϕη*^(*ω*)/*C*^*η*^(*ω*), where 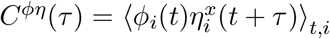 is the cross-covariance of *ϕ*_*i*_(*t*) and the driving noise 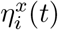 [43].
5. Smoothly update

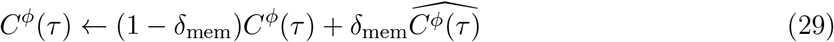

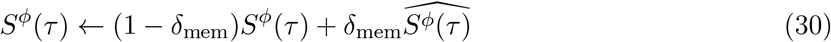

with memory parameter *δ*_*mem*_ ∈ (0, 1).
6. Repeat 2-5 until convergence.

More details for a similar method (for *R* = 1 special case) can be found in [45]. This method was used for the theory fits in Fig. S5B,C,D, with *τ*_*max*_ = 120, *dτ* = 0.05, *N*_*batch*_ = 10^4^, *N*_*iter*_ = 400, and *δ*_*mem*_ = 0.2.

#### Spontaneous state with reservoir of limit cycles

In a special case of an ensemble with constant *D*^(*µ*)^ = *D* and **A**^(*µ*)^ i.i.d. sampled as block-Haar matrices (see Appendix), the single-unit self-coupling kernel *G*^*z*^(*τ*) vanishes. Crucially, this means the currents *x*(*t*) now have Gaussian statistics, and we can leverage closed-form Gaussian integrals in our calculations. Moreover, the kernel that determines input noise to each neuron *Q*^*z*^(*τ*) admits a closed-form expression in Fourier space without the need for an explicit average over tasks *µ*. In turn, *C*^*x*^(*ω*) has a closed-form relationship with *C*^*ϕ*^(*ω*) that is parameterized by the (constant) average gain ⟨*ϕ*′⟩ which depends on *C*^*x*^(*τ* = 0). This can be translated into a second-order ODE (Eq. 150) and solved for as in [36], with the additional step of self-consistently determining the gain.

Alternatively, we can directly solve in the Fourier domain by iteratively enforcing

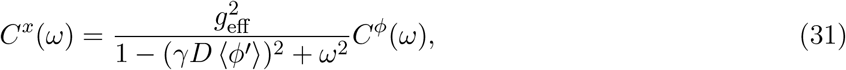

where 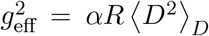 (defined in Eq. (14)), over the whole spectrum. We iterate on the following process:

1. Start with an initial guess for *C*^*x*^(*ω*).
2. Take the inverse Fourier transform to get *C*^*x*^(*τ*).
3. Calculate *C*^*ϕ*^(*τ*) in terms of *C*^*x*^(*τ*) via Eq. (41) and ⟨*ϕ*′⟩ in terms of *C*^*x*^(*τ* = 0) via Eq. (40).
4. Update *C*^*x*^(*ω*) using Eq. (31) with memory parameter *δ*_*mem*_ ∈ (0, 1).
5. Repeat steps 2-4 until convergence.

We evaluated over a temporal window of *τ*_*max*_ = 200, *dτ* = 0.05 with *δ*_*mem*_ = 0.5.

#### Task-selected state with reservoir of limit cycles

For notational simplicity, we assume there is one dominant oscillation, which we index by ∗. We first write down the dynamics for the two latent variables 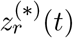 of the dominant pattern with pattern strength *D*^(*∗*)^ and overlap **A**^(*∗*)^, with intrinsic frequency *ω*^(*∗*)^ = tan(*θ*^(*∗*)^). By the assumption of dominance, 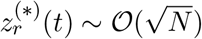, we can divide by a factor of 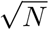, making the 𝒪 (1) noise vanish for large *N*, and get

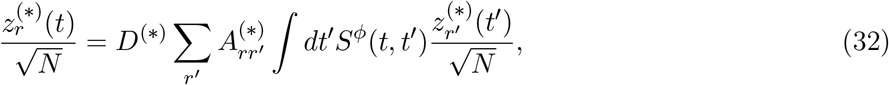

which reduces in the case of Gaussian *x*(*t*) to

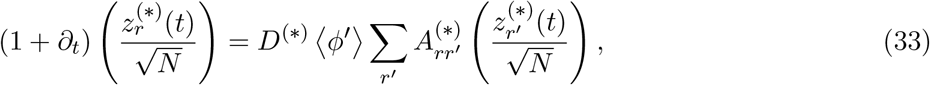

because *S*^*ϕ*^(*t, t*′) = Θ(*t* − *t*′) ⟨*ϕ*′(*t*)⟩ *e*^*−*(*t−t′*)^. These dynamics are also consistent with the direct definition of the condensed patterns from the self-consistency conditions and applying Stein’s lemma to Eq. (26), which the Gaussianity of *x*(*t*) permits. Since *ϕ* is saturating, this system has a steady-state oscillatory solution of a particular amplitude *ρ*. Assuming marginal stability of the oscillation enforces constant ⟨*ϕ*′⟩ = 1/(*γD*^(*∗*)^ cos(*θ*^(*∗*)^), which depends on both *ρ* and the noise variance—though we don’t yet know how this is partitioned between them. For now, we write the steady-state solution for the dominant pattern in terms of *ρ* as

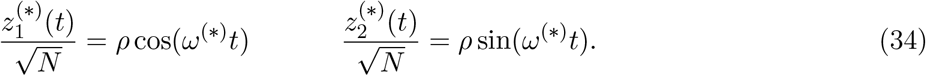

The dynamics for any one neuron *x*_*m*_(*t*) features noise drive from *η*^*x*^(*t*) sampled iid as a Gaussian process with stationary kernel *Q*^*z*^(*τ*), determined like before as an average over the 2-point functions of the weak-task trajectories, plus deterministic drive from this dominant task with coupling *m*^(*∗,r*)^:

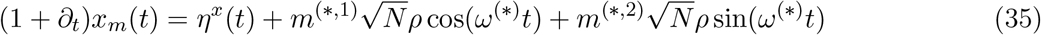

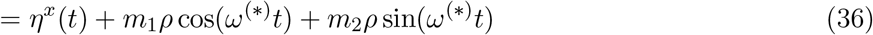

where we have defined 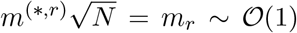 as the order-1 set of couplings between this arbitrary unit and the condensed oscillation, which encode both amplitude and relative phase information. The *P* subscript in *x*_*m*_(*t*) indicates this dependence on 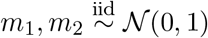.

As currently setup, this problem is reminiscent of [37], but with a few key differences:

- The noise is not due to an i.i.d. reservoir, but rather a reservoir of low-rank matrices with 2 × 2-block Haar overlaps. This has an effect on the temporal shape of the noise statistics (the difference between Eq. 150 and the equation for an unstructured network).
- In [37], neural inputs feature phase heterogeneity across units but uniform amplitude. Our case has heterogeneity in both amplitude and phase. This changes how we write down our average *ϕ* autocovariance function.
- The overall scaling of the (heterogenenous) amplitudes *ρ*∥*m*∥ is not a free parameter but rather self-consistently determined with *D*^(*∗*)^, the constant *D* on the uncondensed patterns, and current variance. This changes the axes on the bifurcation diagram, since *D*^(*∗*)^ is what we freely vary and not *ρ*, but once equilibrium *ρ* is determined, the effect on the dynamics is similar to that of the external current from [37].

We write 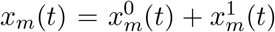 as a sum of a filtered deterministic part and a filtered noise part, the former having a closed-form expression:

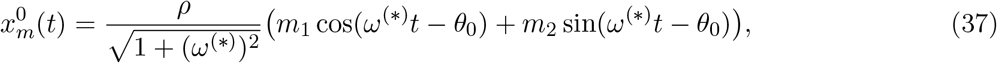

which has autocovariance 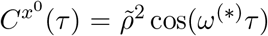 for 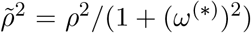. The filtered noise part *x*^1^(*t*) has autocovariance with the same relation to *C*^*ϕ*^(*ω*) as in the spontaneous case:

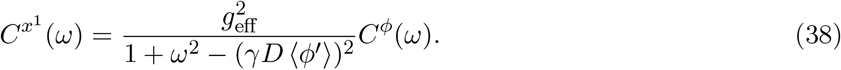

(See Appendix for a derivation of this relation.) If we can write an expression for *C*^*ϕ*^(*τ*) in terms of the statistics of 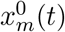 and 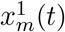, then we close the equations. Since they are both Gaussian, their sum is Gaussian, and 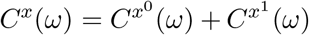 determines *C*^*ϕ*^(*τ*) in closed form (after taking the IFFT). That suggests the following iterative process for solving the DMFT:

1. For a given *D*^(*∗*)^ and *ω*^(*∗*)^, first calculate the average gain ⟨*ϕ*′⟩ set by the marginality condition of the dominant oscillation.
2. Set an initial guess 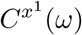.
3. Take the IFFT to obtain 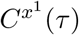.
4. Invert Eq. (40) to solve for the complementary 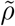 value that combines with 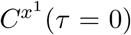 to give the already determined ⟨*ϕ*′⟩ value.
5. Compute 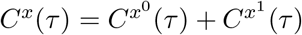 in terms of our provisional 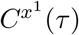 and 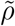 values.
6. Compute *C*^*ϕ*^(*τ*) using Eq. (41).
7. Take the FFT of *C*^*ϕ*^(*τ*) to obtain *C*^*ϕ*^(*ω*)
8. Update estimate for 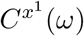 using *C*^*ϕ*^(*ω*) via Eq. (38) with memory parameter *δ*_*mem*_ ∈ (0, 1).
9. Repeat steps 2-8 until convergence.

We evaluated over a temporal window of *τ*_*max*_ = 200, *dτ* = 0.05 with *δ*_*mem*_ = 0.5. Additionally, for a given dominant frequency *ω*^(*∗*)^ value, we adjusted the value to precisely match the nearest on-bin frequency value to avoid spectral smearing.

#### Phase boundaries

Computing 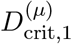 is straightforward from Eq. (21) and only requires solving the spontaneous state DMFT and computing ⟨*ϕ*′⟩. To compute 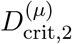, we used a midpoint solver to identify the value of 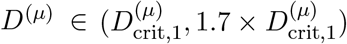 that results in a *C*^*x*^(*τ*) curve with

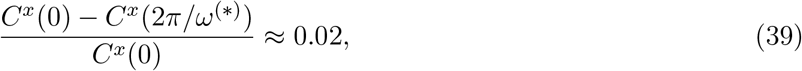

where 0.02 is our criterion for when the relative magnitude of fluctuations is effectively vanished.

#### Gaussian integrals

Any special case with Gaussian currents (e.g. the reservoir of oscillations) can take advantage of Gaussian integrals to simplify calculation. By our choice of nonlinearity 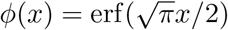, we can leverage certain closed-form expressions:

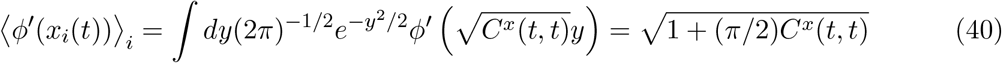

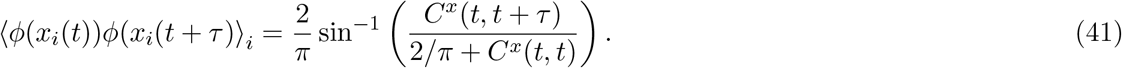

Whenever we consider Gaussian currents *x*_*i*_(*t*) in this work, they are always zero mean with possibly time-dependent variance *C*^*x*^(*t, t*), but more often we are in the stationary case *τ* = *t* −*t*′ with *C*^*x*^(0) = *C*^*x*^(*t, t*).

### Dimensionality of task-selected state

See [45] for a detailed walkthrough of dimensionality calculations, which can in principle apply to the spontaneous (𝒪 (*N*) dimensionality) or task-selected (𝒪 (1) dimensionality) states. However, in practice we found poor numerical fits for dimensionality in the spontaneous state when overlaps were even modestly strong.

The dimensionality calculation simplifies tremendously for the task-selected state. We can leverage the duality between neuron-neuron covariance (used most directly to define the PR) and time-time covariance (which has identical spectrum to the neuron-neuron covariance) as first shown in [45]. For a given variable *a* ∈ *{x, ϕ}*, its PR can be computed in terms of its autocovariance function *C*^*a*^(*τ*) and four-point function 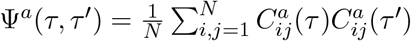

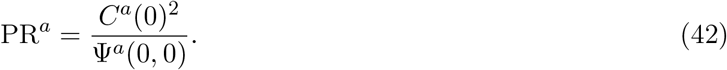

Leveraging the duality discussed above, we can alternatively calculate Ψ^*a*^(*τ, τ*′) as

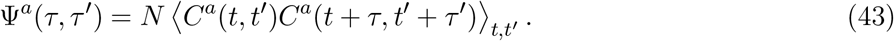

For *τ* = *τ*′ = 0, we’re essentially computing the average square of *C*^*a*^(*t, t*′) over the whole (*t, t*′) plane. The values near the *t* = *t*′ diagonal contribute negligibly compared to the whole plane, so we need only examine the asymptotic off-diagonal behavior when *t*−*t*′ is large. Any fluctuations *δC*^*a*^(*t, t*′) = *C*^*a*^(*t, t*′)−*C*^*a*^(*t*−*t*′) about the stationarized two-point function are negligible in this case, since typical values over the plane are 𝒪 (1) because of the dominant dynamics. Thus we can estimate Ψ^*a*^(0, 0) directly from the autocovariance function *C*^*a*^(*τ*) as

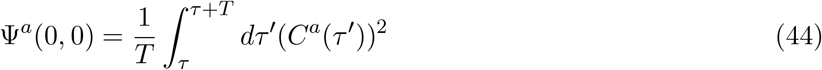

where the domain of integration depends on the nature of the dominant task. For a limit-cycle task, we need only integrate over one period of an oscillation to get the average behavior over the whole plane, as long as we start past when the amplitude decay is finished (Fig. S7). For a fixed point, we need only integrate over a brief interval when the *C*^*a*^(*τ*) has reached its asymptotic value.

## Simulating networks

To validate the theoretical calculations, we simulated networks as well. We varied *N* depending on how much we needed to reduce finite-size effects in a given context, we always used *dt* = 0.05 and RK4 forward integration. Parameters are in Table 1 for all network simulations in the main and supplementary figures.

**Table 1:** Summary of network parameters used in simulations across figures. In all cases, *R* = 2.

| Panels | $N$ | $P$ or $\alpha$ | $D^{(\mu)}$ | Notes on $A^{(\mu)}$ / sampling |
| --- | --- | --- | --- | --- |
| Fig. 2A | 2000 | $P = 1$ | $D = 2.2$ | $\mathbf{A} = \begin{pmatrix} 0.8 & 0.4 \\ -0.4 & 0.8 \end{pmatrix}$ (limit cycle) |
| Fig. 2B | 2000 | $P = 1$ | $D = 2$ | $\mathbf{A} = \begin{pmatrix} 0.5 & 0.3 \\ 0.3 & 0.5 \end{pmatrix}$ (bistable) |
| Fig. 2C | 2000 | $P = 2$ | $D^{(1)} = 2.2$, $D^{(2)} = 2$ | $\mathbf{A}^{(1)}$ limit cycle, and $\mathbf{A}^{(2)}$ bistable |
| Fig. 2D,E,F | varied | varied | $D^{(1)} = 3$, $D^{(\mu>1)} = 2$ | $\mathbf{A}^{(1)}$ limit cycle; $\mathbf{A}^{(2)}$ bistable; $A_{rr'}^{(\mu>2)}$ sampled i.i.d. $\mathcal{N}(0, 0.5)$ with rejection of non-positive-semidefinite draws |
| Fig. 2G | 4000 | $\alpha = 0.5$ | $D^{(1)} = 2$ or 9; $D^{(\mu>1)} = 2$ | $\mathbf{A}^{(1)}$ limit cycle; $\mathbf{A}^{(2)}$ bistable; $A_{rr'}^{(\mu>2)}$ sampled i.i.d. $\mathcal{N}(0, 0.6)$ with rejection of non-positive-semidefinite draws |
| Fig. 3C,D,E | 5000 | $\alpha = 0.5$ | $D^{(\mu)} = 2$ by default; $D^{(1)}$ modulated from $2 \rightarrow 3.75 \rightarrow 8$; $D^{(2)}$ from $2 \rightarrow 4.2 \rightarrow 8$ | $\mathbf{A}^{(1)}$ limit cycle; $\mathbf{A}^{(2)}$ bistable; $A_{rr'}^{(\mu>2)}$ sampled i.i.d. $\mathcal{N}(0, 0.6)$ with rejection of non-positive-semidefinite draws |
| Fig. 4A | 4000 | varied | $D^{(\nu \neq \mu)} = 2$, and $D^{(\mu)}$ varied, where $\mu$ corresponds to limit cycle of frequency $\omega = 0.25$. | $\mathbf{A}^{(\nu \neq \mu)} = \gamma \begin{pmatrix} \cos \theta & \sin \theta \\ -\sin \theta & \cos \theta \end{pmatrix}$ with $\theta \sim \text{Unif}(0, 2\pi)$ (see Appendix) |
| Fig. 4B | 4000 | $\alpha = 1$ | $D^{(\mu)}$ varied on y-axis, and $\mu$ varied on x-axis by as the intrinsic frequency $\omega = \tan \theta$ for $\mathbf{A}^{(\mu)}$. | same as above |
| Fig. 5A,B,C | 8000 | $\alpha = 1$ | same $D^{(\mu)}$ as Fig. 4 modulated from $2 \rightarrow 4.15 \rightarrow 5.3$ (black boxes), | same as Fig. 4 |
| Fig. 5D | varied | $\alpha = 1$ | same $D^{(\mu)}$ modulated | same as Fig. 4 |
| Fig. 5E | varied | $\alpha = 1$ | same as Fig. 4 | |
| Fig. S2 | 2000 | $P = 2$ | $D^{(1)} = 2$, $D^{(2)} = 2.2$ | same as Fig. 2C |
| Fig. S3A | 2000 | $P = 2$ | $D^{(1)} = 3.5$, $D^{(2)} = 4$ (or swapped) | same as Fig. 2C |
| Fig. S3B | 2000 | $P = 2$ | $D^{(1)} = 3.3$, $D^{(2)} = 3$ (or swapped) | same as Fig. 2C |
| Fig. S4 | 5000 | $P = 2$ | $D^{(\mu)} = 2$ or $2 \rightarrow 2.75 \rightarrow 3.5 \rightarrow 4.25$ | same as Fig. 3 |
| Fig. S5A-D | 8000 | $P = 2$ | $D^{(\mu)} = 2$ | same as Fig. 3 |
| Fig. S6 | 10000 | $P = 2$ | $D^{(\nu \neq \mu)} = 2 \rightarrow 4 \rightarrow 8$ | same as Fig. 4 |

### Software availability

Core functions for simulating networks and solving the theory are available in https://github.com/omarschall/mft-theory, as well as an example notebook showcasing the fits in Fig. 5C.

## Additional mathematical details

### Connectivity and connection to SVD

Eq. (4) can be equivalently interpreted as the product of three matrices: an *N* ×*PR* matrix whose columns comprise the left loading vectors **m**^(*µ,r*)^, an *PR* × *PR* diagonal matrix whose diagonal elements comprise the task weights *D*^(*µ*)^ each repeated *R* times, and an *PR* × *N* matrix whose rows comprise the right loading vectors **n**^(*µ,r*)^ (Fig. S1A). Thus the multi-task network (Eq. 4) is of the same form as a low-rank network with rank *PR* (Eq. 2).

Just as the Singular Value Decomposition (SVD) decomposes a matrix as a weighted sum of outer products, our model *generates* a matrix as a weighted sum of outer products (Fig. S1A). The loadings **m**^(*µ,r*)^ (**n**^(*µ,r*)^) play the role of the left (right) singular vectors, except they are not perfectly orthonormal but rather orthonormal up to 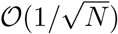 noise, i.e. 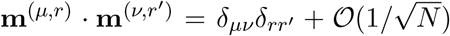. The unrolled index *a* = 1, …, *M* ×*R* corresponding to (*µ, r*) lists the “singular values” *D*^(*a*)^ for each of the *P* ×*R* modes (Fig. S1B). For a general matrix, these could be any non-negative values, but here we constrain them to be constant within *R* blocks, i.e. *D*^(*µ,r*)^ = *D*^(*µ,r′*)^≡ *D*^(*µ*)^. For an arbitrary matrix, the overlaps between left and right singular vectors are generally unconstrained up to basic geometric consistency, and similarly we are free to generate connectivity through a nearly arbitrary pattern of overlaps *O*_*ab*_ = ⟨ **n**^(*a*)^ *·* **m**^(*b*)^ ⟩ (Fig. S1C). We consider the special case of overlap matrices with an *R* × *R* block-diagonal structure (Fig. S1C), because keeping *R* fixed and allowing *P* → ∞ makes the model mathematically tractable. However, the set of overlaps as specified by 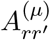 is arbitrary^3^. This formulation is equivalent to the “random-mode model” with overlaps from [45], except the symmetries have a block-diagonal, rather than merely diagonal (equivalently *R* = 1), structure.

### Full details of small-*P* case

The small-*P* case is equivalent to the single-task network case, because if *P* is small then *R* × *P* is also small. In the large-*N* limit, there is zero direct cross-talk between latent variables from distinct tasks, and Eq. (3) factorizes into self-contained systems of size *R*

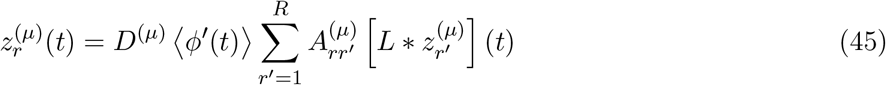

up to their sharing of the gain factor via the Gaussian integral of *ϕ*′ with variance given by the squared norm of all the low-pass-filtered latent variables 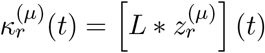:

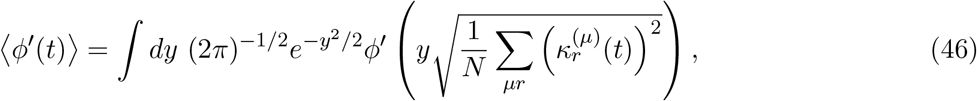

which can be evaluated by Eq. (40). Here *L*(*τ*) = Θ(*τ*)*e*^*−τ*^ is a low-pass filter, a closed-form special case of the filter *F*(*τ*) that we can leverage with *G*(*τ*) = 0, as in particular occurs when *α* = 0 (i.e. small *P*) but in other cases as well.

### Winner-take-all intuition

We present intuition for why there must always be a dominant task in this case. For a task *µ* in this setting to be dominant, its linearized dynamics must be unstable about the origin, so that it grows until 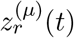 is large enough to influence the nonlinearity ⟨*ϕ*′(*t*)⟩ and self-stabilize. (This necessarily requires 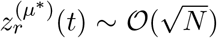 because of the *N* scaling in the argument of *ϕ*′.) The dynamics are marginally stable when

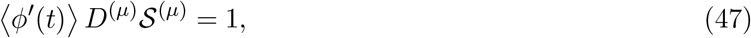

where 𝒮^(*µ*)^ = max[ℜ 𝔢 (*λ*) |*λ* eigenvalue of **A**^(*µ*)^]. If no task even achieves *D*^(*µ*)^ *𝒮*^(*µ*)^ ≥ 1 with maximal gain ⟨*ϕ*′(*t*)⟩ = 1, then the network is in the quiescent state, with all activity decaying to the origin rapidly. To avoid this trivial case, we assume at least one task is sufficiently strong to avoid the quiescent state. Consider the “strongest” such task, i.e. the one with highest value of *D*^(*µ*)^ 𝒮^(*µ*)^. To keep itself balanced, this task must set ⟨*ϕ*′(*t*)⟩ ∼ *D*^(*µ*)^ 𝒮^(*µ*)^ so that its dynamics are either marginally stable, or alternate phases of growth and decay cyclically (e.g. an elliptical limit cycle). Since all other tasks share this gain factor and have by definition weaker intrinsic strength, each will have a strictly weaker recurrent drive and therefore feature contracting dynamics. This is true even in cases where multiple tasks are fine-tuned to have the same strength—finite-sampling effects will always result in one task being marginally stronger than another. However, the more fine-tuned two tasks are to have the same strength, the more likely particular network realizations can feature long transient periods of activity where tasks appear “co-dominant,” until one eventually takes over.

What makes Gaussian loadings in particular lead to only one task dominating, is the fact that, in such networks, there is only one “population” in the sense of [29]. Thus there is only one gain factor, which equilibrates to a value that marginally stabilizes the strongest dynamics and suppresses all weaker ones, as described above. In a network with multiple populations, multiple gain factors could in principle separately equilibrate to values that marginally stabilize two different tasks’ dynamics—a trivial example would be adjoining two completely unconnected single-task networks that we call one network. However, nontrivial examples with overlap and interaction among populations also exist, which we leave to future work.

### Fixed-point task dominance

In the main text, we showed an example where the limit cycle dynamics dominate the fixed-point dynamics. Here we swap the *D*^(*µ*)^ values such that the bistable dynamics dominate, and the activity in the task-1 subspace decays to the origin (Fig. S2A,B). The two matrices generating the limit-cycle and fixed-point dynamics, respectively,

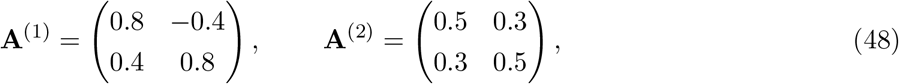

were chosen such that they have identical spectral abscissa of *S*^(1)^ = *S*^(2)^ = 0.8, so that the dominant task is whichever simply has larger *D*^(*µ*)^.

## Supplemental Figures and Tables

**Figure S1:**
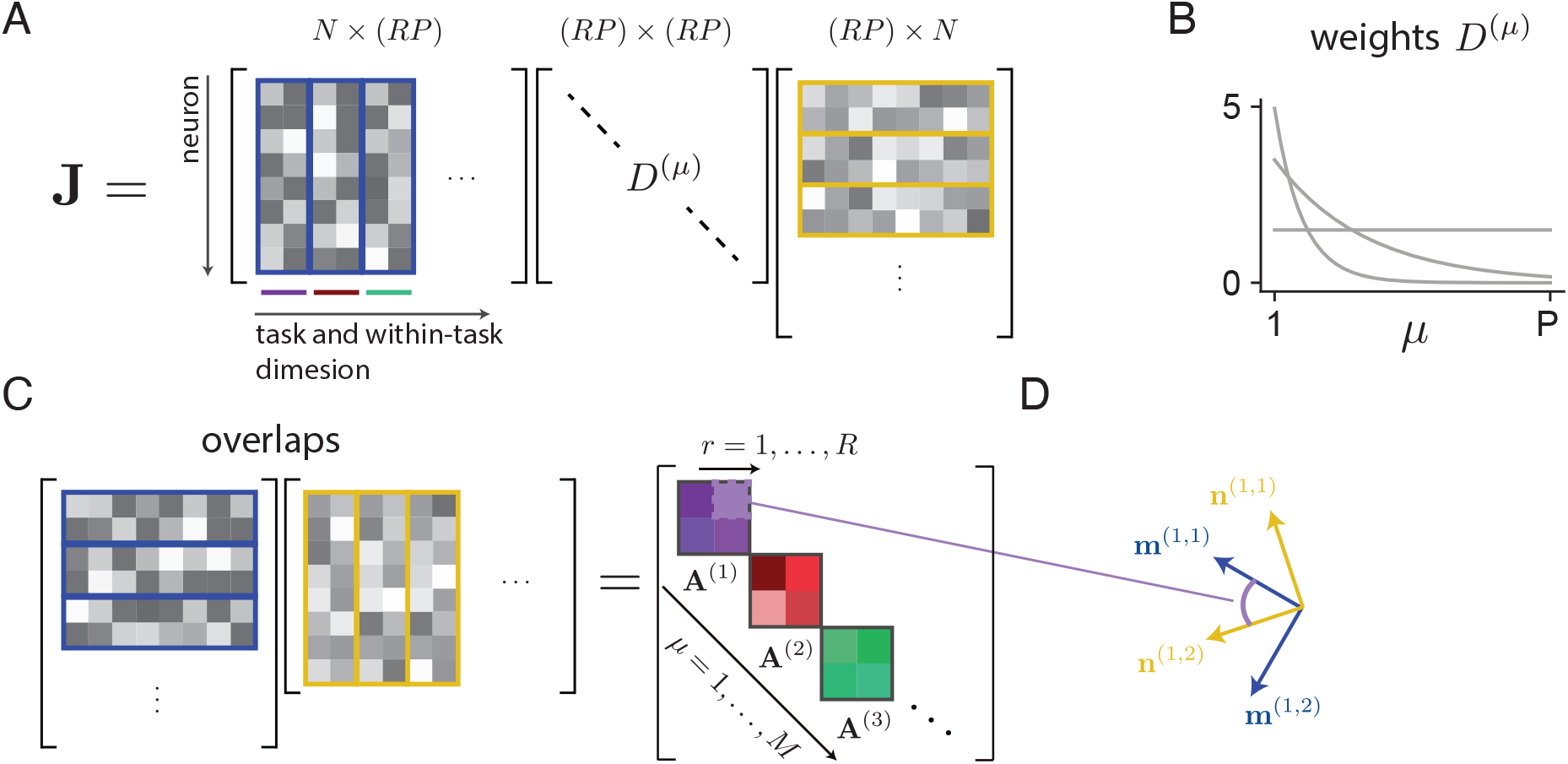
More detailed illustrations of connectivity structure. **(A)** Alternative interpretation of the sum in Fig. 1C as one large matrix product, with left loadings organized in one matrix as columns (blue) and right loadings in another as rows (yellow), with scalings as diagonal elements. The blue or yellow rectangular boundaries group the loading vectors into tasks, so *r* = 1, 2 indexes vectors within a block, while *µ* = 1, …, *P* indexes rectangular blocks. **(B)** Example connectivity strength spectra *D*^(*µ*)^, which can generally be dispersed or concentrated among tasks. **(C)** Illustration of the overall overlap matrix for all loading vectors, grouped by task by blue or yellow rectangular boundaries, as the matrix product of the left loading vectors as rows of one matrix (blue) with the right loading vectors as columns of one matrix (yellow). **(D)** Geometric interpretation of an entry of the overlap matrix as the degree of alignment of the associated left and right loading vectors. The **m**’s (and **n**’s) are shown to be orthogonal among themselves.

**Figure S2:**
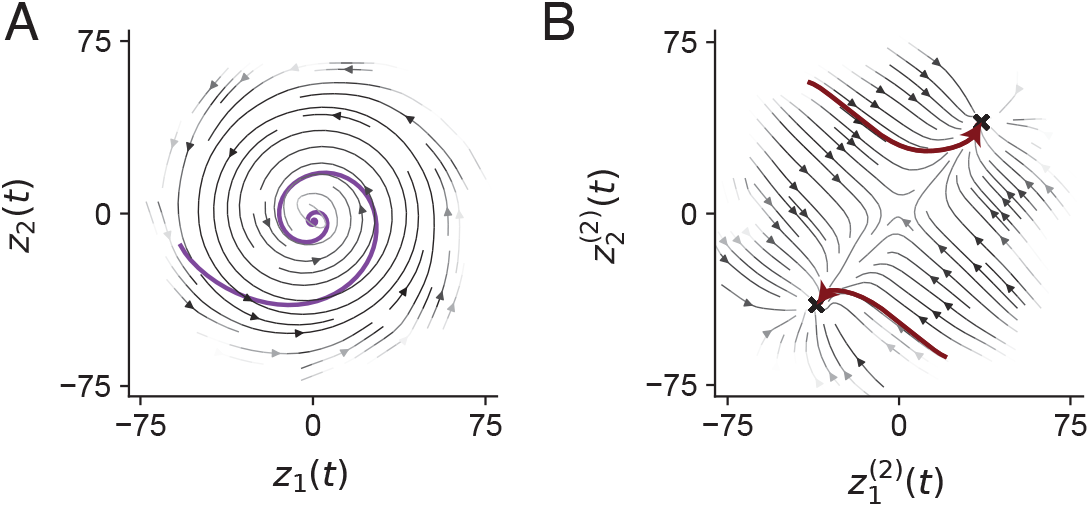
Dynamics for the same network as Fig. 2C but with *D*^(1)^ = 2, *D*^(2)^ = 2.2. **(A)** Purple: activity in the task-1 subspace for a single run of the dynamics. Black: flow fields over many runs. **(B)** Red: activity in the task-2 subspace over two runs with different initial conditions. Black: average flow fields over many runs.

**Figure S3:**
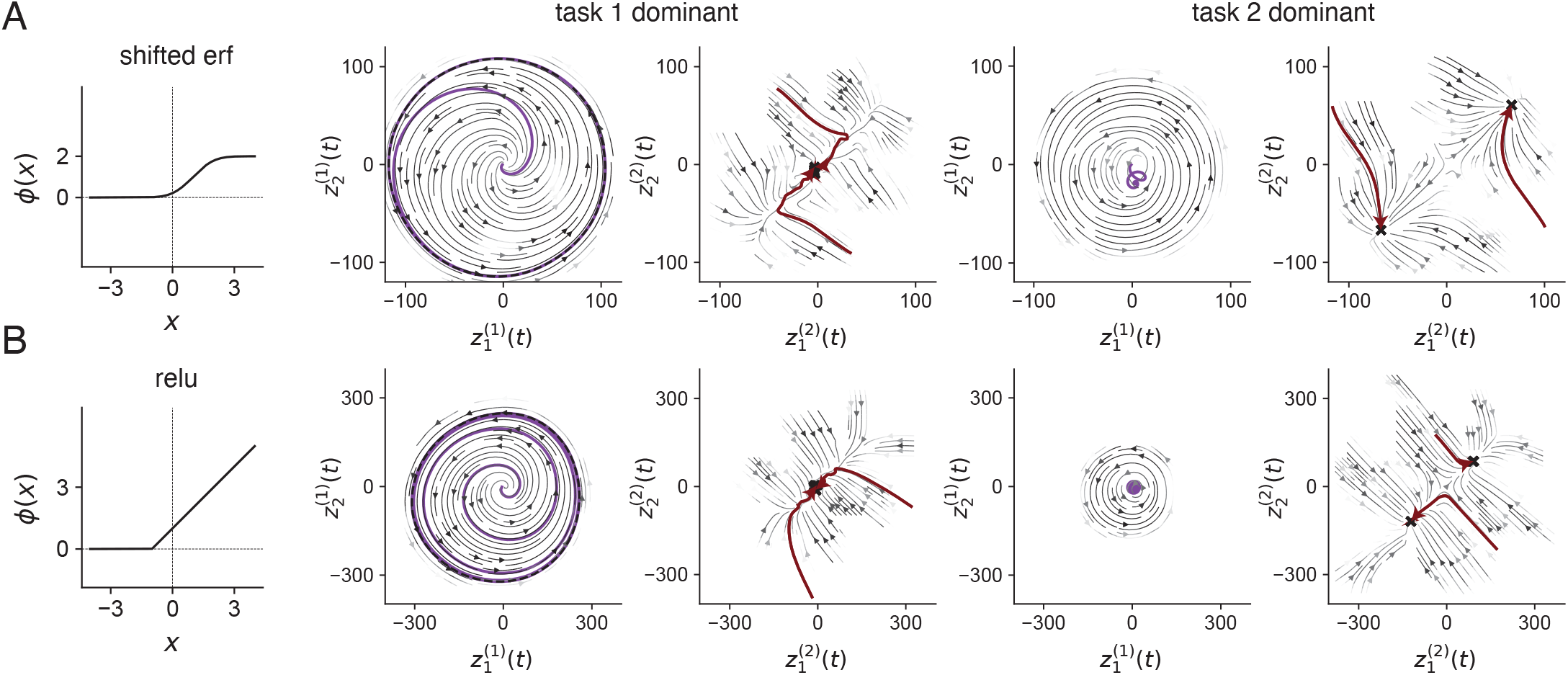
Similar simulations as in Fig. 2, but for networks with different nonlinearities. **(A)** A network using a diagonally shifted error function, by 1 in each direction, such that outputs are always positive, and it requires positive current to activate a neuron. **(B)** A network using a non-saturating nonlinearity, in particular a relu function (shifted leftward by 1). To ensure global stability, an additional term −*g*_*I*_/*N* is added to each *J*_*ij*_, with *g*_*I*_ = 2.

**Figure S4:**
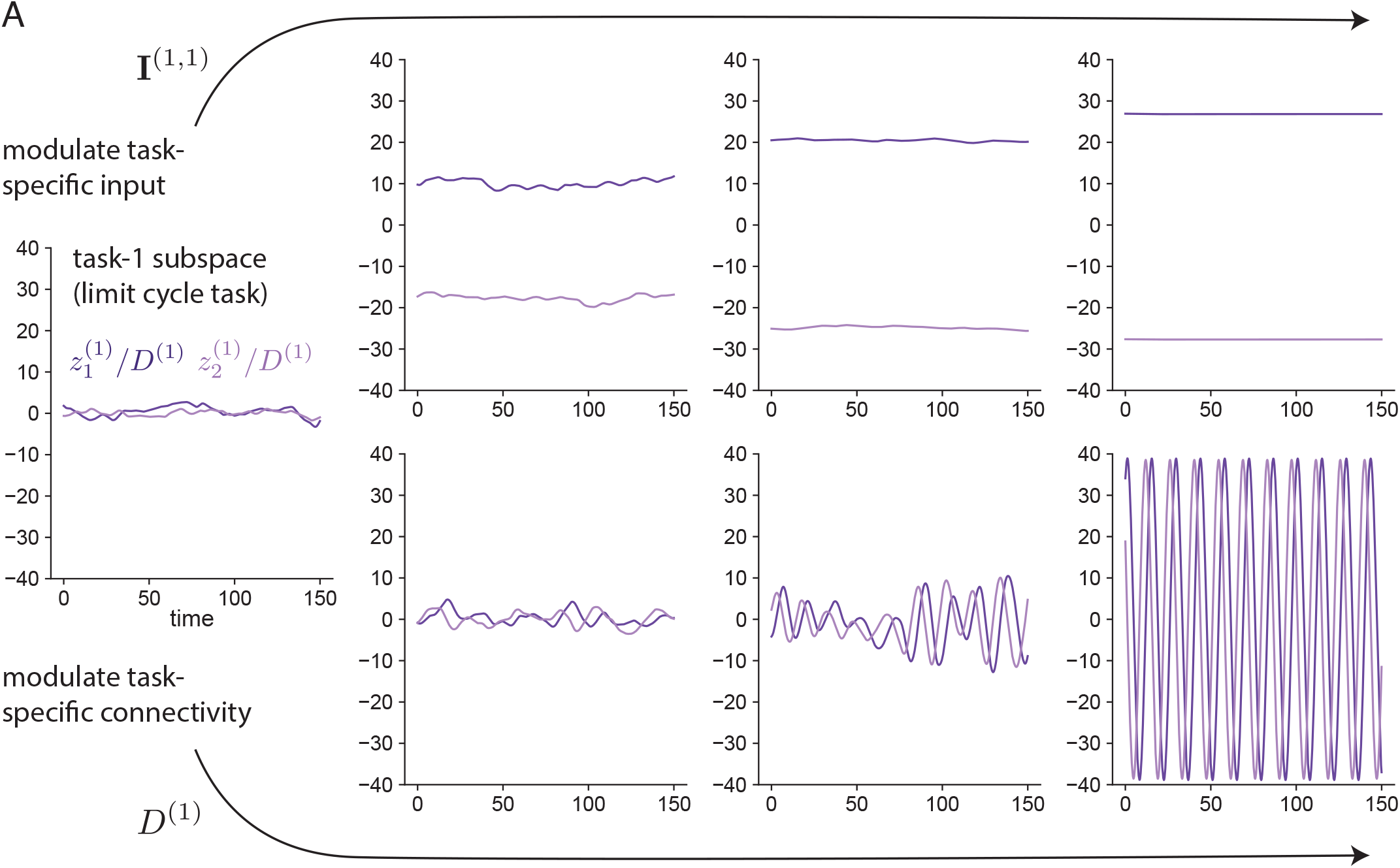
Comparing modulation of task-specific input drive and task-specific connectivity, for task *µ* = 1 in the same network shown in Fig. 3. **(A)** Left: spontaneous-state activity in the task-1 subspace, featuring weak, noisy oscillations. Top: modulating the additive input drive 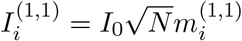 to each neuron *i* in the direction of the first task-1 left loading vector. Values of *I*_0_ are (left to right) 0, 0.5, 1, 1.5. Bottom: modulating the strength of the task-1 connectivity component. Values of *D*^(1)^ are (left to right) 2, 2.75, 3.5, 4.25.

**Figure S5:**
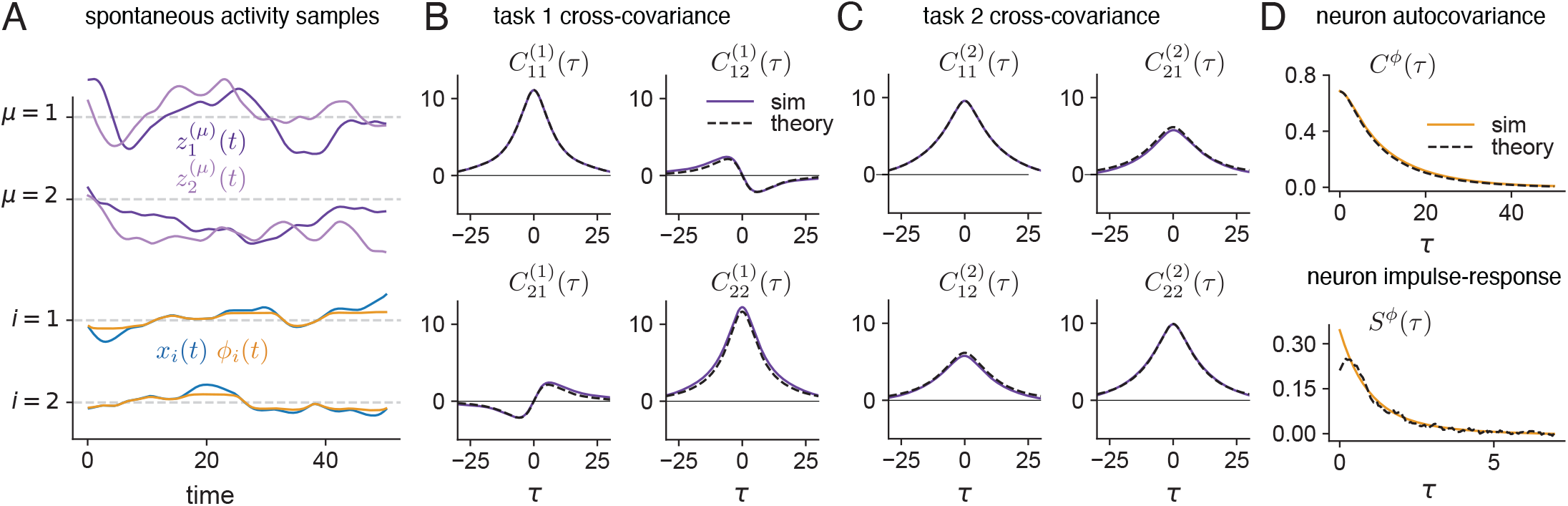
Dynamics in the spontaneous state. **(A)** Example activity traces for a network with *α* = 0.5; *D*^(*µ*)^ = 2 for all *µ*; **A**^(1)^, **A**^(2)^ as in Fig. 2C, with 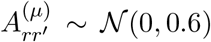 i.i.d. with rejection sampling for positive-definiteness violations; and *N* = 8000. Top: latent activity for a limit-cycle task (*µ* = 1) and a fixed-point task (*µ* = 2) in the spontaneous state. Bottom: current and firing-rate traces for two example neurons in the network. **(B)** Lagged cross-covariance functions for 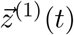, i.e. 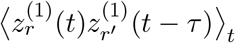. Top left: *r* = 1, *r*′ = 1. Top right: *r* = 1, *r*′ = 2, etc. Solid: simulation. Dashed: theory. **(D)** Single-neuron autocovariance function *C*^*ϕ*^(*τ*) = ⟨ *ϕ*_*i*_(*t*)*ϕ*_*i*_(*t* + *τ*) ⟩_*t*_ (top) and impulse-response function *S*^*ϕ*^(*τ*) = ⟨*ϕ*′⟩ *F*(*τ*) (bottom) and for the same network. Solid: simulation. Dashed: theory.

**Figure S6:**
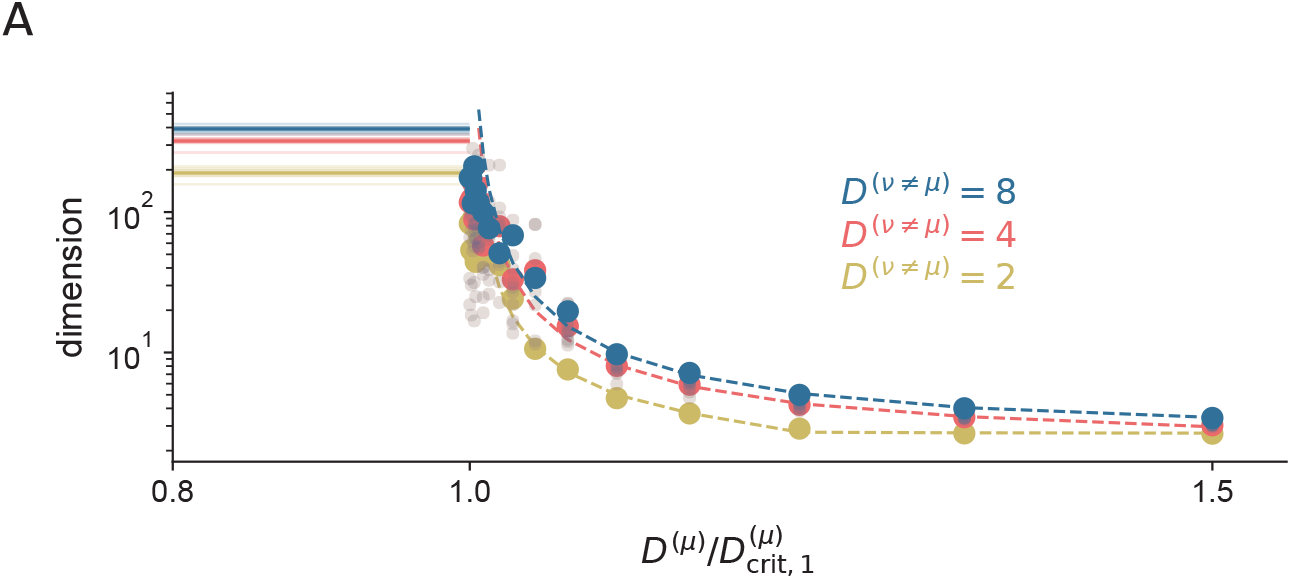
**(A)** Participation ratio of firing rates for networks with variable *D*^(*ν*)^ values for all non-selected tasks ν ≠ *µ*, as *D*^(*µ*)^ is varied relative to its critical point for task selection. Light dots: individual network simulations. Heavy dots: averages over networks. Dashed line: theory.

**Figure S7:**
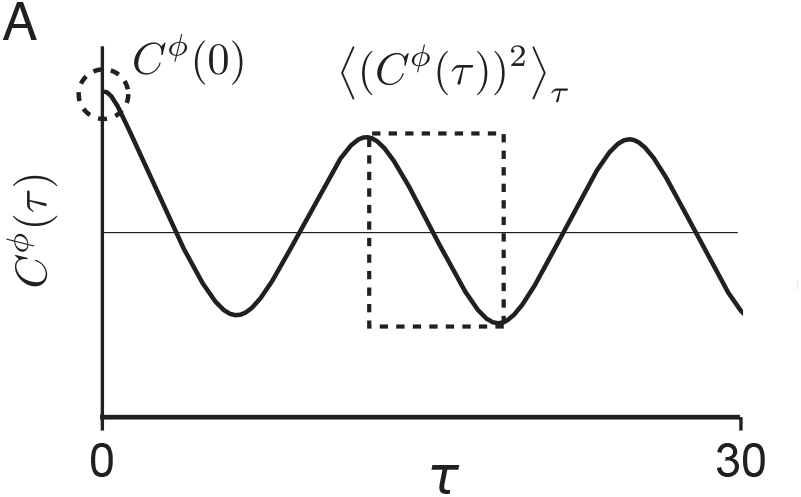
Cartoon showing how to read off the dimensionality of the task-selected state from the single-unit autocovariance function. **(A)** Black: Theoretically calculated autocovariance function *C*^*ϕ*^(*τ*) from Fig. 3F (middle). The value at *τ* = 0 at the average squared value over one period after full amplitude decay are the relevant quantities for computing the PR. The latter captures asymptotic behavior of the autocovariance function away from *τ* = 0, which is needed for Eq. (44).

## Appendix: Derivations of key mathematical results

### Path integral for spontaneous state

This section takes advantage of mathematical techniques which are outlined in e.g. [68] or [69]. Similar derivations can be found in [45]. In this section, we derive the DMFT for the spontaneous state, assuming no dominant patterns. We start by writing the Martin-Siggia-Rose-Janssen-de Dominicis path integral for the partition function *Z*(**J**) of the network dynamics:

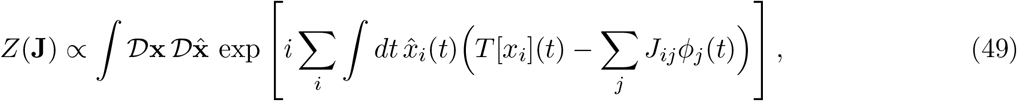

currently keeping the weights as shorthand *J*_*ij*_ and using *T* = 1 + ∂_*t*_ as notational convenience. We now compute the average

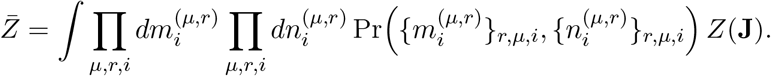

The probability density Pr(· · ·) for a given set of loadings^4^ for each neuron *i* and each task *µ*,

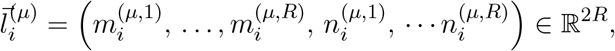

is

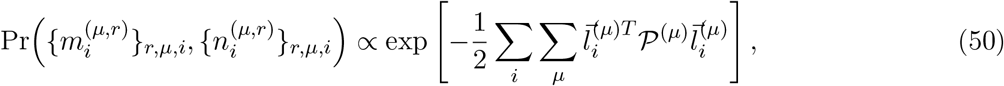

since each neuron’s loadings are independent of the others’, and loadings for different tasks are also independent. Within a task for a given neuron, the loadings are distributed as a multivariate Gaussian with precision matrix

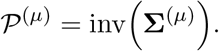

We define a set of *R* × *P* latent variables

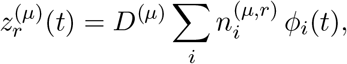

and their definition-enforcing conjugates 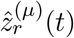. Then our disorder-averaged partition function is

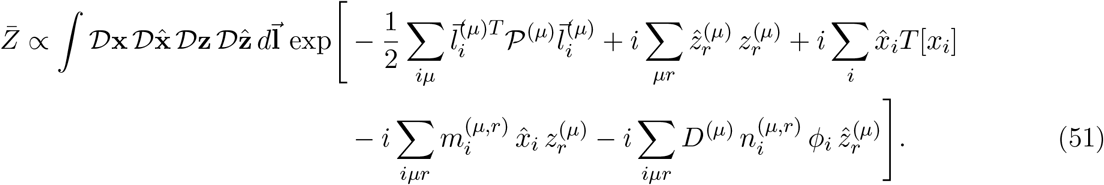

where suppressed time indices imply time integrals over a shared dummy time index. We can rewrite this as a quadratic in terms of 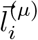 by defining the vector

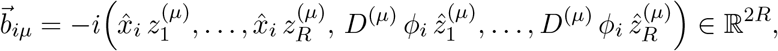

allowing us to perform a matrix version of completing the square for each *iµ*:

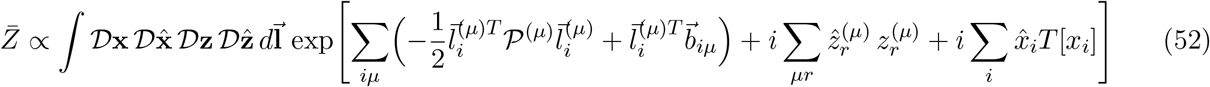

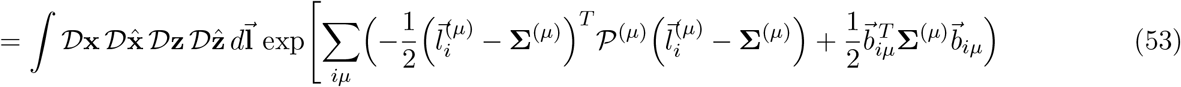

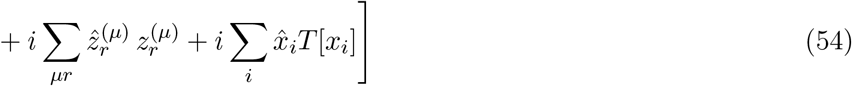

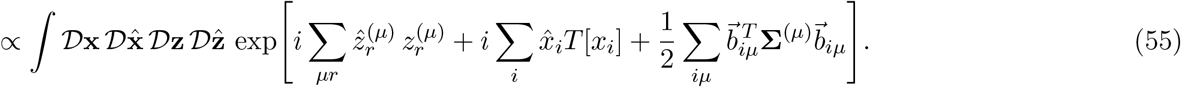

Recall that **∑**^(*µ*)^ has a block diagonal structure with the identity along each of the two *R* × *R* principal blocks. Then we can rewrite this quadratic term as three terms, reintroducing the time indices briefly:

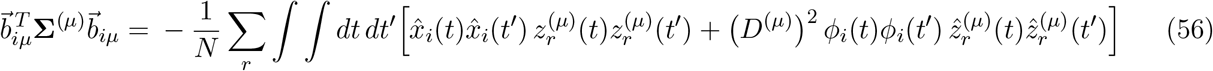

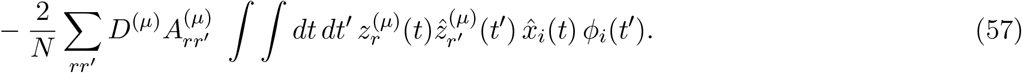

The first two terms are easy to deal with and imply a reciprocal relationship between the two-point function of the latent variables 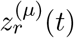 and how it appears in the noise driving *x*_*i*_(*t*), and the two-point function of the units *ϕ*_*i*_(*t*) and how it appears in the noise driving 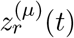. In the random-mode-model without symmetry [45], these are the only terms, and they lead to a straightforward solution. The third term, which arises due to the latent weight symmetries encoded in the off-diagonal block 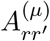 of **∑**^(*µ*)^, is more difficult to deal with. We have to split the expression into the *t* > *t*′ and *t* < *t*′ half-planes. In the former, we can write 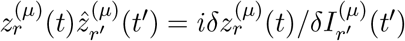, and in the latter we can write 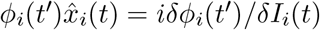, where *δ*/*δI*_*i*_(*t*) represents the functional derivative with respect to external current. (In the flipped cases, each is zero by causality.)

Then we define a set of fields and their conjugates:

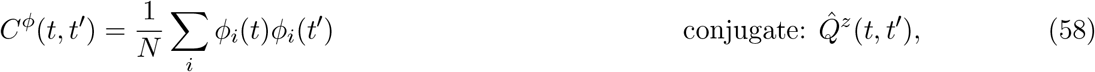

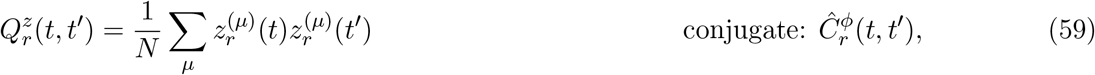

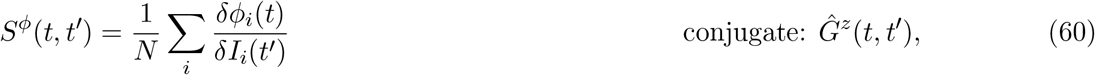

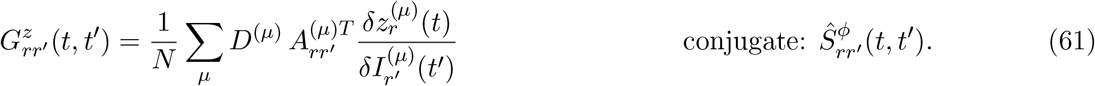

Plugging in these definitions, we have

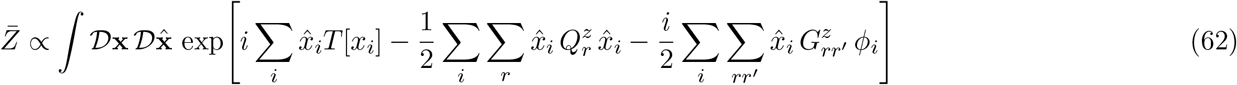

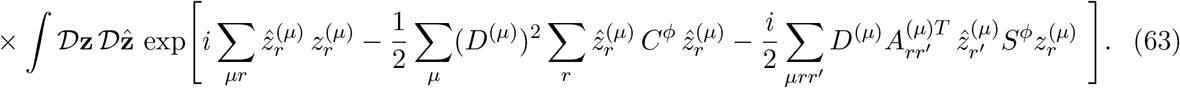

Note how the interaction term has a half in the neuron path integral and a half in the latent path integral, with kernels defined as population-averaged susceptibilities in the other picture. In other words, a given neuron interacts with itself by a weighted average of latent-latent impulse responses, and a given set of same-task latent variables interact among themselves by an average of single-neuron impulse responses.

Then, integrating over the definitions of these fields and decoupling noninteracting single-neuron and single-task path integrals, we obtain definition-enforcing terms and a global N scaling outside an action 1066 that is purely a function of the fields:

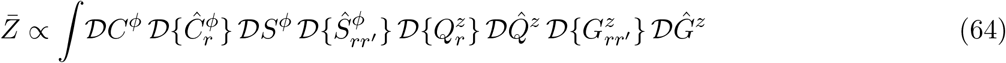

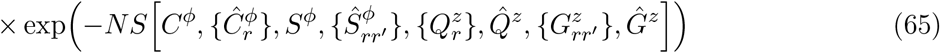

with

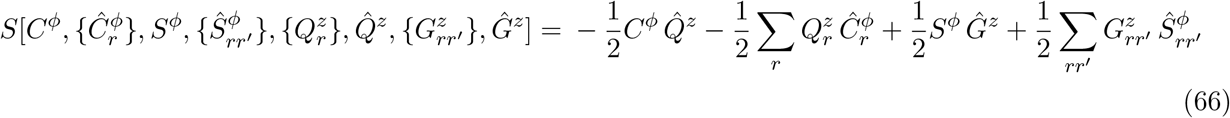

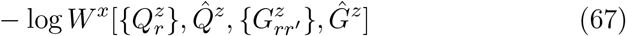

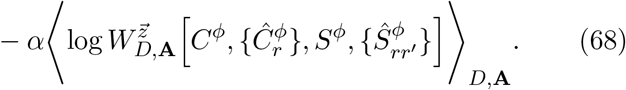

Here,

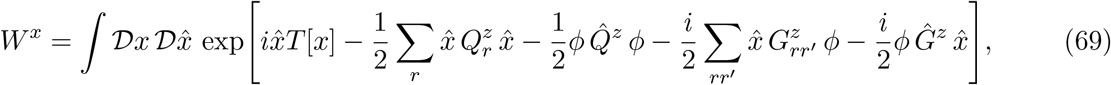

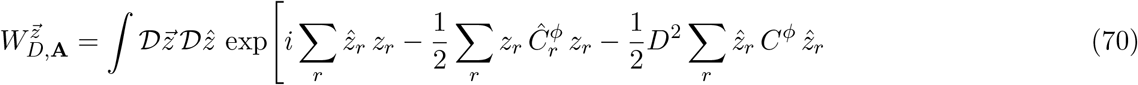

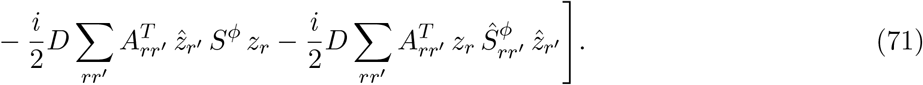

We differentiate with respect to each of the *R*^2^ + *R* + 2 fields and their conjugates:

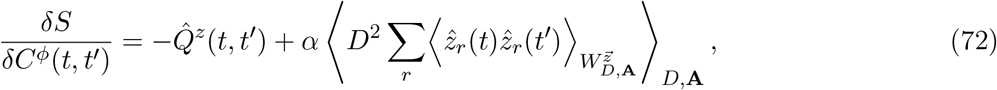

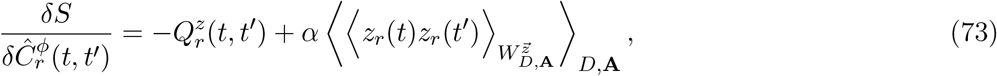

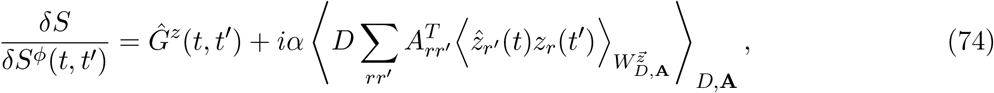

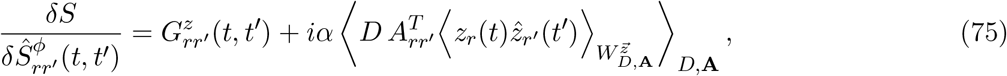

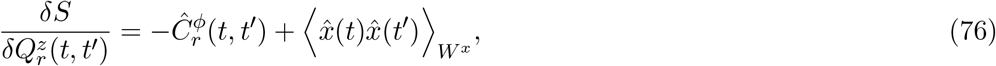

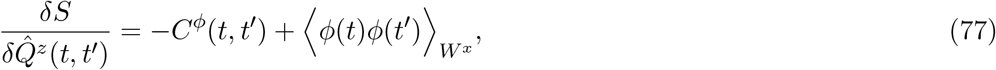

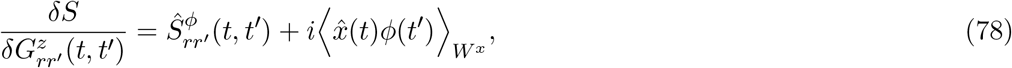

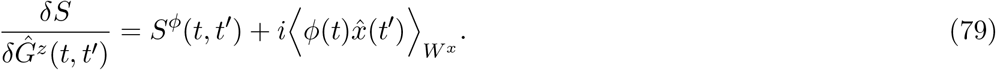

Solving the saddle-point system yields

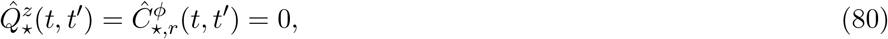

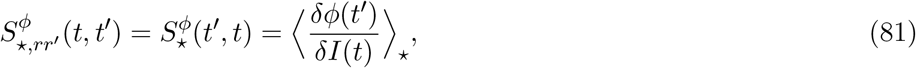

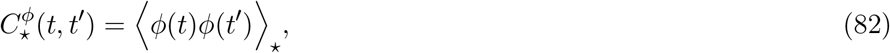

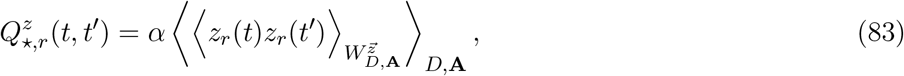

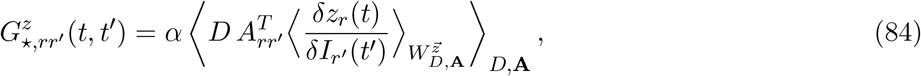

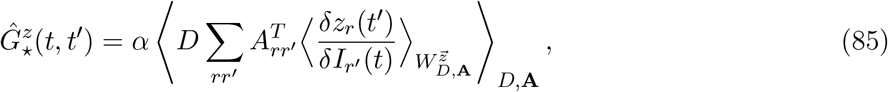

so that

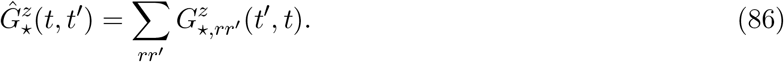

Then, going back into the single-site and latent processes, we obtain a *ϕ* process driven by noise term 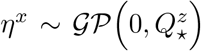 where 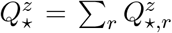, plus self-interactions via self-coupling kernel 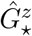. However, the process for the latent variables is more complex and involves *R*^2^ self-couplings. Altogether, we have

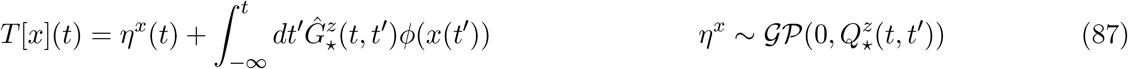

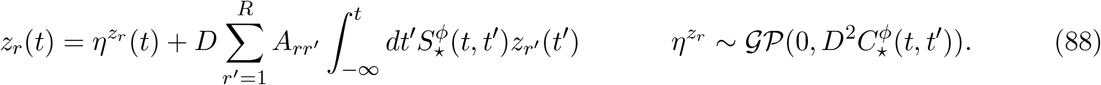

From now on, we drop the ⋆ notation and assume any kernel as written is the saddle-point solution. Further, we assume that all kernels depend only on *t*−*t*′, allowing us to leverage the convolution theorem. In Fourier space, we solve for *z*_*r*_(*ω*) and get

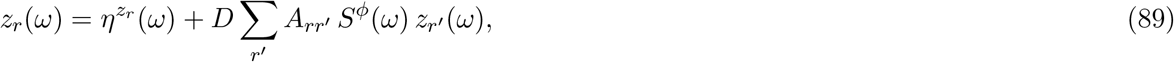

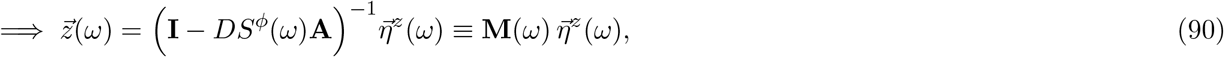

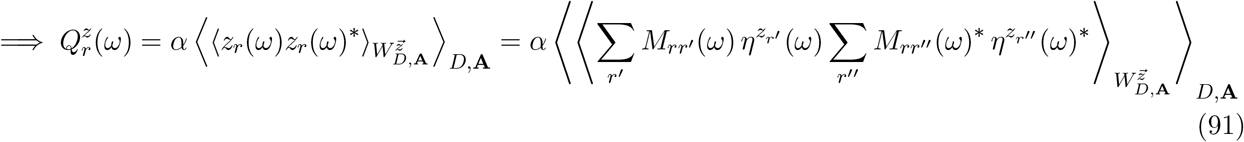

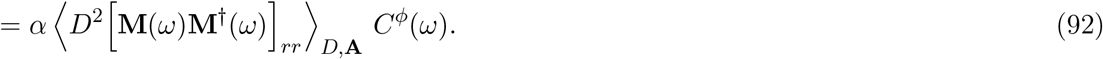

Then using Furutsu–Novikov we get an expression for 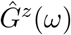. We summarize together with *Q*^*z*^(*ω*) as

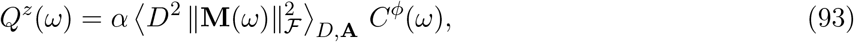

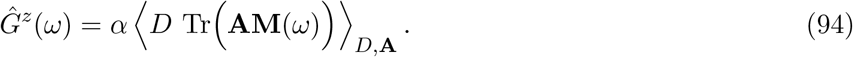

### Path integral for task-selected state

We pick up at Eq. (55) and explicitly separate the *µ* index into a set of *P*^*∗*^ dominant and *P* − *P*^*∗*^ ≈ *P* weak tasks. We perform the disorder integration only over the weak tasks’ couplings and keep the average partition function depending on the particular realization of the dominant couplings:

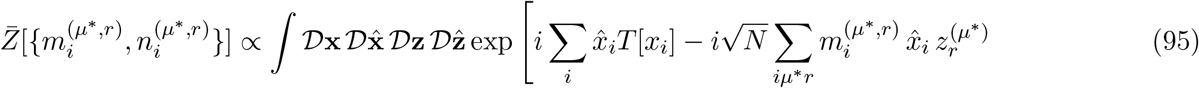

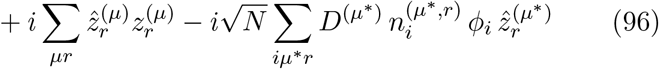

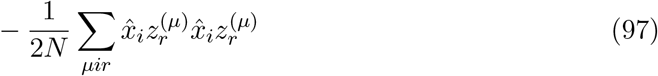

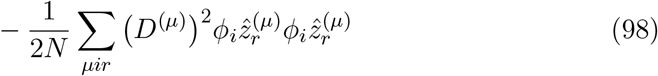

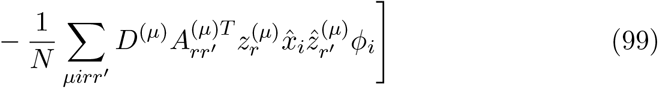

We now define an expanded set of order parameters

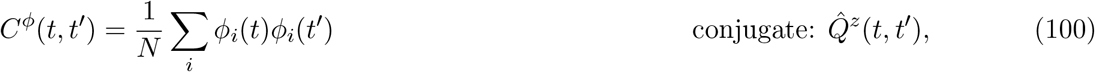

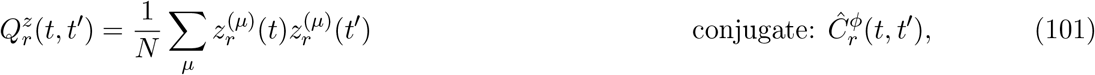

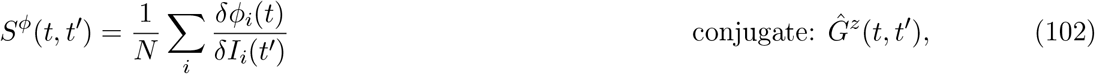

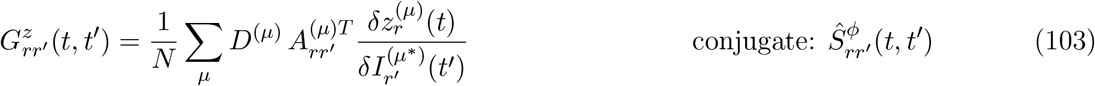

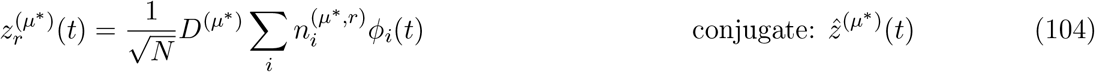

For the dominant tasks, their activity is no longer a dynamical or stochastic variable but rather a trajectory that is subject to the saddle-point condition of the overall dynamics. Note that we have rescaled the usual definition of the task latents by a factor of 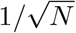 so that all order parameters are 𝒪 (1). We insert definitions of order parameters when possible and separate into integrals over **x**’s and **z**’s.

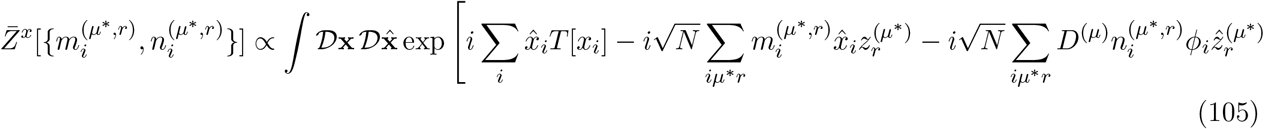

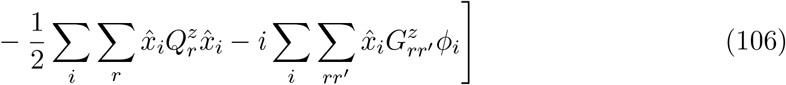

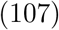

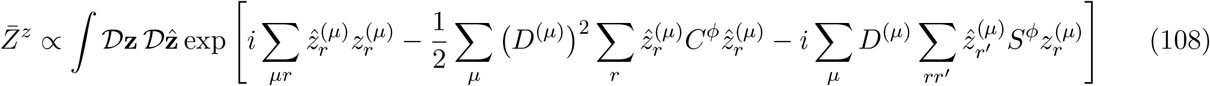

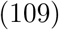

Then we can construct one path integral over the fields as a function of the realizations of the normalized dominant couplings 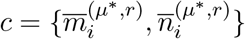:

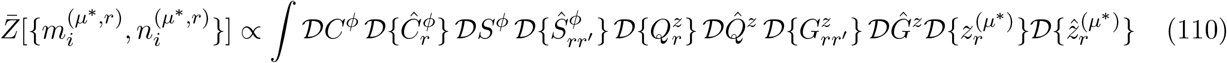

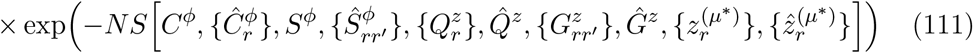

with

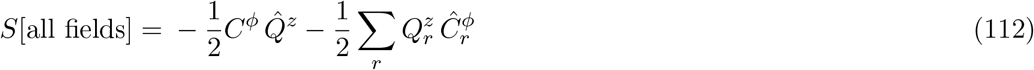

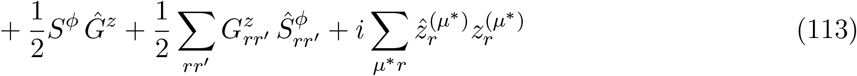

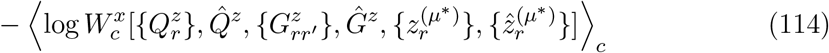

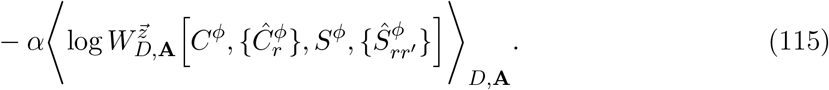

Here,

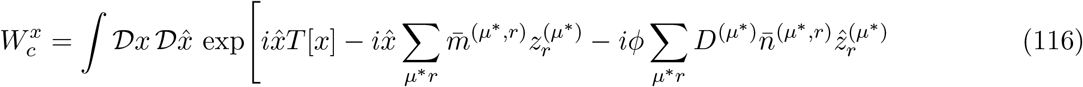

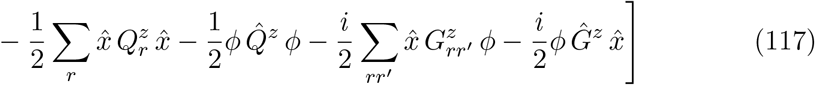

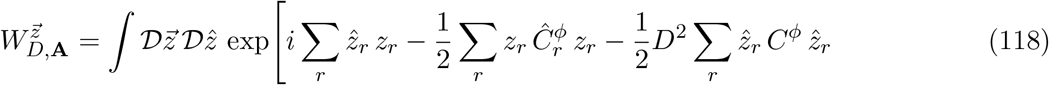

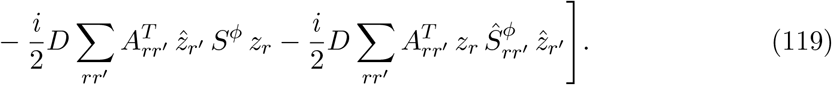

where we have defined 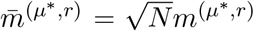 (and similarly 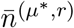) as the 𝒪 (1) normalized couplings of an arbitrary unit to the dominant patterns. Then differentiating with respect to each order parameter, we get

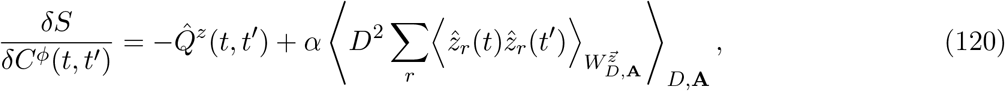

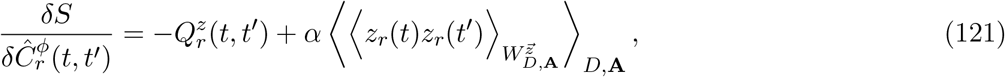

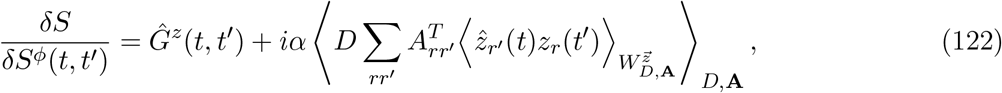

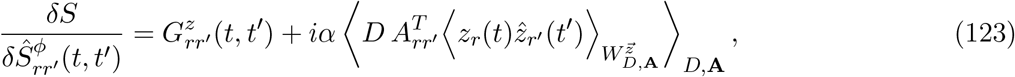

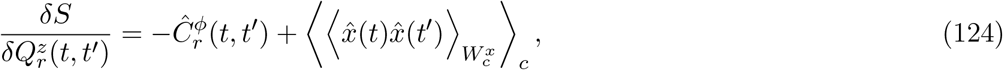

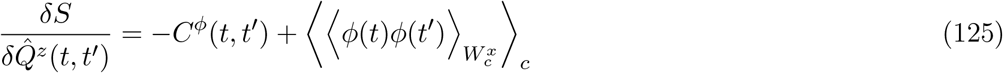

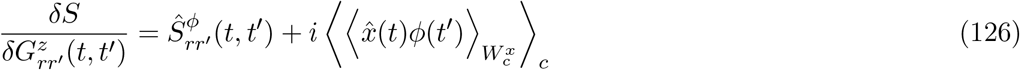

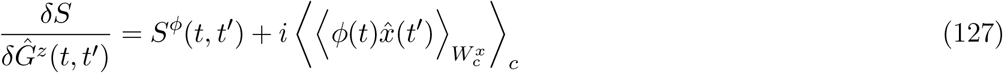

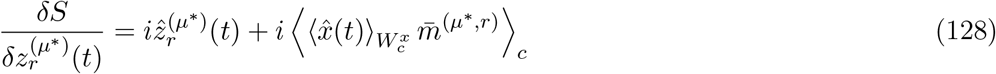

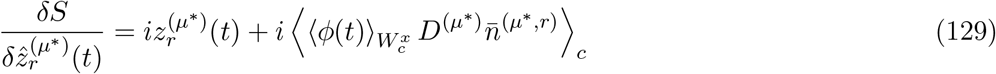

We get the same self-consistency conditions as before, except now the single-site properties are averages over heterogeneous units, where the particular sample of 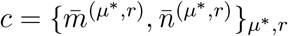 for that unit determines its dynamics and properties, plus two additional conditions:

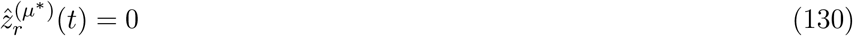

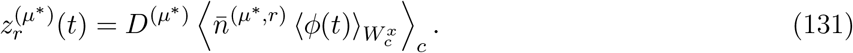

Then we have our final mean-field theory

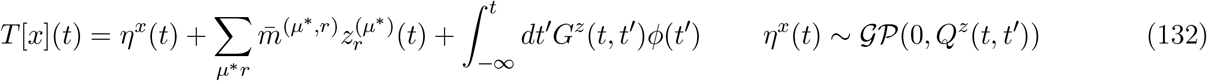

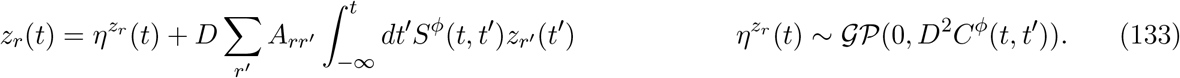

In the main text a factor of 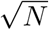 gets traded between 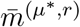 and the non-normalized dominant pattern 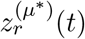.

### DMFT simplifications for ensemble of limit-cycle tasks

Consider the following process for generating *R*×*R* matrices, where *R* must be even. We start by sampling *R*/2 independent 2 × 2 rotation matrices matrices via *θ*_*l*_ ∼ *U*(0, 2*π*) for l = 1, …, R/2:

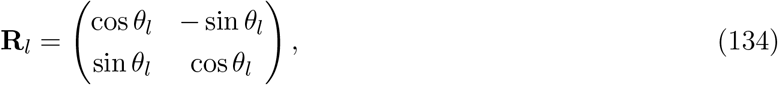

and arrange them in a block-diagonal structure. We then take a random *R* × *R* Haar orthogonal matrix **B** and intermix the entries, but preserving the traces of all matrix powers. We finally scale the result by *γ* < 1 (e.g. *γ* = 0.99) to ensure that 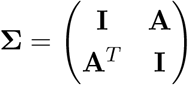 is positive definite. Then

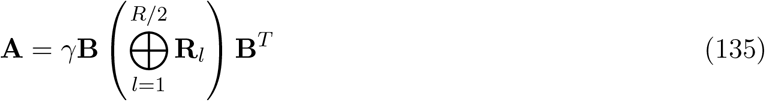

has a few desirable properties as a process for randomly generating a matrix. (For simplicity, the main text considers just *R* = 2, **B** = **I** =⇒ **A** = *γ***R**_1_, although its useful properties extend to this more general case.) First, for any *k* ≥ 1, we have

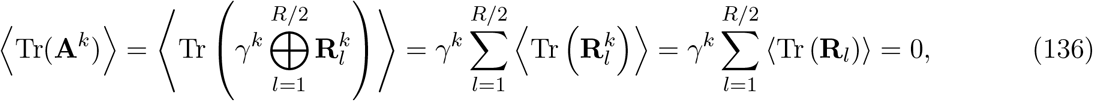

using the fact that 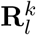 has the same distribution as **R**_*l*_. The *γ* factor conveniently also reduces the SEM when we effectively “measure” the mean trace matrix power for finite samples of such matrices in simulated networks, which can be large for large powers *k*.

For independently sampled *D*, this property eliminates the single-unit self-coupling by setting *G*^*z*^(*t, t*′) = 0, but without resorting to a trivial distribution of **A** = **0** as is necessary and sufficient in the *R* = 1 case. We also have a closed-form equation for *Q*^*z*^(*ω*), which reads

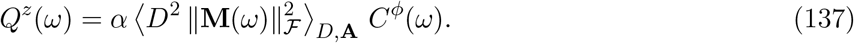

where **M**(*ω*) = (**I** −*DS*^*ϕ*^(*ω*)**A**)^*−*1^. Under these assumptions about the distribution of **A**, we can calculate *Q*^*z*^(*ω*). First, we note that the similarity transformation pops out of the entire **M**(*ω*) matrix:

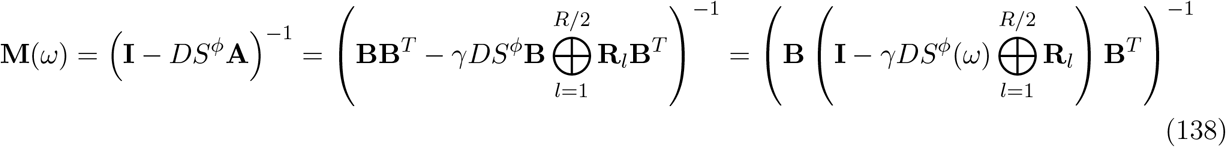

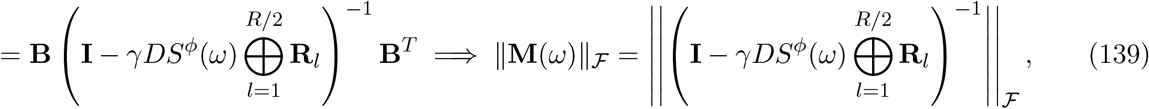

since the Frobenius norm is invariant under orthogonal similarity transformations. Next we apply the matrix inversion on each block separately:

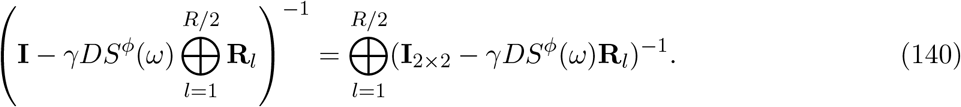

Each component matrix that gets inverted **I**_2*×*2_ − *γDS*^*ϕ*^(*ω*)**R**_*l*_ has eigenvalues 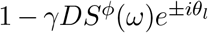, so their inverses have eigenvalues 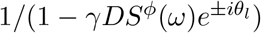. Since this matrix is clearly normal, its Frobenius norm squared is simply the sum of the eigenvalues’ squared norms. Putting it all together and assuming *D* is sampled independently from **A**, we have

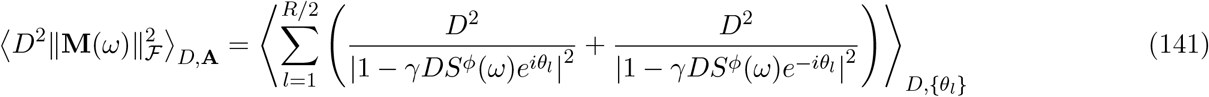

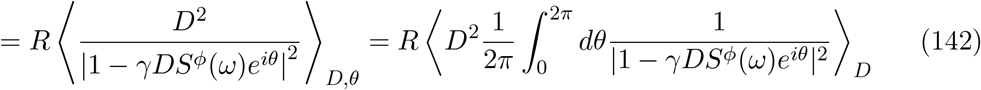

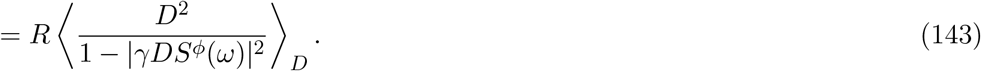

The last line requires assuming |*γDS*^*ϕ*^(*ω*)| < 1, which is a fairly safe ass
umption for a distribution of *D* with no outliers. Then, in Fourier space, the statistics of the effective neuron noise follow

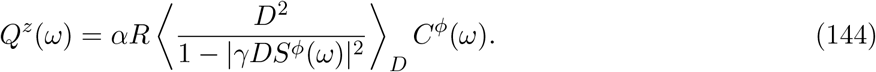

With these simplifications, we have now a bipartite mean-field theory

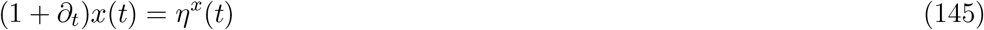

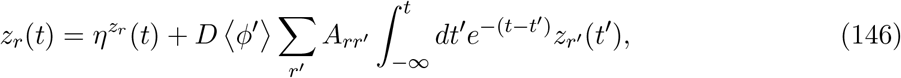

because we know *S*^*ϕ*^(*τ*) = Θ(*τ*) ⟨*ϕ*′⟩ *e*^*−τ*^ and *G*^*z*^(*τ*) = 0. We can differentiate the second equation and get

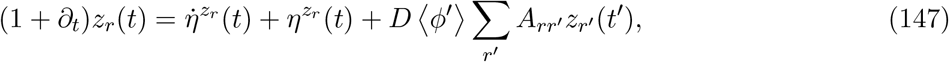

exploiting the fact that the derivative of *S*^*ϕ*^(*τ*) is negative of itself, producing a leak term. With this representation of the weak-task latent variable dynamics, it is much more straightforward to determine the stability condition on *D*: it reduces to whether the largest real part among eigenvalues of *D* ⟨*ϕ*′⟩ **A** is below or above 1.

Let’s examine the single-site picture and solve for the temporal dynamics of the *x*(*t*) autocovariance:

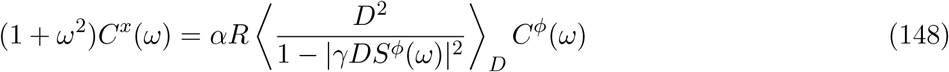

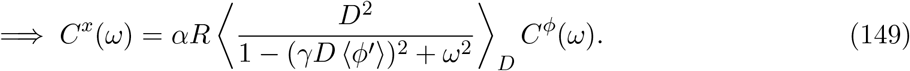

Now let’s make yet a further assumption, that *D* is in fact a constant such that 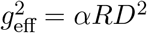 rather than following some distribution—this is a common choice for the non-selected tasks. Then we have

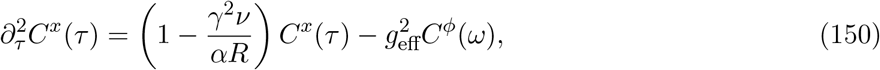

which is exactly the second-order ODE describing *C*^*x*^(*τ*) for an unstructured network [36], except the *C*^*x*^(*τ*) term is scaled by 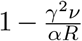 of 1, where ν ≡ (*g*_*eff*_ ⟨*ϕ*′⟩)^2^ can be thought of as the effective slope for an equivalent linear network [43, 70].

The average gain depends exclusively on *C*^*x*^(0) (via a Gaussian integral). However, to solve for *C*^*x*^(0) by the Sompolinsky potential-energy trick in this case requires knowing the average gain already, because of its appearance in *ν*. Therefore we solve for *C*^*x*^(0) for a provisional value of ⟨*ϕ*′⟩, which we iteratively update as we solve for *C*^*x*^(0) again with our new value of ⟨*ϕ*′⟩, until it stops changing. This value of ⟨*ϕ*′⟩ then determines the stability for an arbitrary task, based on the spectral abscissa of *D* ⟨*ϕ*′⟩ **A**. However, the purely spectral methods we describe earlier are more effective for small *α*.

## Footnotes

1 Although in the main text we emphasized that there can be at most one dominant task, this is true for sufficiently long times (to forget initial conditions) and when there are no external inputs. Generally, initial conditions or external input drive can set multiple task-related latent variables to be large. Once inputs are released, the network eventually equilibrates to having at most one dominant task. In all cases, only finitely many tasks can be dominant since the state vector norm is at most 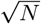.

2 With a small difference: ⟨*ϕ*′(*t*)⟩ is determined by 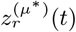 as well as a constant interference term from non-dominant tasks. But since this term has Gaussian statistics, Stein’s Lemma still applies.

3 Up to the constraint that the overall 2*R ×* 2*R* loadings covariance matrix, containing the *n*-*n* block 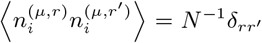 and *m*-*m* block 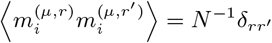 *′* as well as the *n*-*m* blocks given by 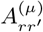, must be positive definite.

4 In general, vector-arrow notation imply a fixed-dimension vector like 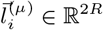, while bolded notation implies a vector with extensive (i.e. *N*-scaling) dimension, e.g. **m**^(*µ,r*)^ ∈ ℝ^*N*^.

